# Hemopexin Reverses Activation of Lung eIF2a and Decreases Mitochondrial Injury in Chlorine Exposed Mice

**DOI:** 10.1101/2023.08.17.553717

**Authors:** Sadis Matalon, Zhihong Yu, Shubham Dubey, Israr Ahmad, Emily M. Stephens, Ammar Saadoon Alishlash, Ashley Meyers, Douglas Cossar, Donald Stewart, Edward P. Acosta, Kyoko Kojima, Tamas Jilling, James A. Mobley

## Abstract

We assessed the mechanisms by which non-encapsulated heme, released in the plasma of mice post exposure to chlorine (Cl_2_) gas, resulted in the initiation and propagation of acute lung injury. We exposed adult C57BL/6 male and female to Cl_2_ (500 ppm for 30 min) in environmental chambers and returned them to room air and injected them intramuscularly with a single dose of human hemopexin (hHPX; 5 µg/ g BW), the most efficient scavenger of heme, 30-60 min post exposure. Concentrations of hHPX in plasma of air and Cl_2_ exposed mice were 9081±900 vs. 1879± 293 at 6 h and 2966±463 vs. 1555±250 at 50 h post injection (ng/ml; X±1 SEM=3; p<0.01). Cl_2_ exposed mice developed progressive acute lung injury post exposure characterized by increased concentrations of plasma heme, marked inflammatory response, respiratory acidosis and increased concentrations of plasma proteins in the alveolar space. Injection of hHPX decreased the onset of acute lung injury at 24 h post exposure; mean survival, for the saline and hHPX groups were 40 vs. 80% (P<0.001) at 15 d post exposure. Non-supervised global proteomics analysis of mouse lungs at 24 h post exposure, revealed the upregulation of 92 and downregulation of 145 lung proteins. Injection of hHPX at one h post exposure moderated the Cl_2_ induced changes in eighty-three of these 237 lung proteins. System biology analysis of the global proteomics data showed that hHPX reversed changes in mitochondrial dysfunction and elF2 and integrin signaling. Western blot analysis of lung tissue showed significant increase of phosphorylated elF2 at 24 h post exposure in vehicle treated mice but normal levels in those injected with hHPX. Similarly, RT-PCR analysis of lung tissue showed that hHPX reversed the onset of mtDNA lesions. A form of recombinant human hemopexin generated in tobacco plants was equally effective in reversing acute lung and mtDNA injury. The results of this study offer new insights as to the mechanisms by which exposure to Cl_2_ results in acute lung injury and to the therapeutic effects of hemopexin.

## Introduction

When humans are exposed accidentally to chlorine Cl_2_ released in the atmosphere because of industrial accidents, they experience varying degrees of acute pulmonary and systemic injury, which may lead to death from respiratory and multi-organ failure. Survivors may develop airway hyper-responsiveness (a prelude to asthma), peri-bronchial and interstitial fibrosis and pulmonary emphysema (2, 3). Medical treatment with supplemental oxygen, bronchodilators and mechanical ventilation are supportive and no specific countermeasures exist.

Cell-free heme (CFH) is an important source of hydroxyl radicals via the Fenton reaction (33), of alkoxyl and peroxyl radicals via conversion of organic hydroperoxides (34) and intercalates and disrupts cell membranes (17). A preponderance of evidence shows that CFH is pro-oxidant, pro-inflammatory and it has been implicated in several pathologies such as malaria, sepsis and exacerbations of sickle cell disease, to name a few(8, 24, 36). For this reason, CFH is maintained at a low level via multiple redundancies served by heme binding proteins such as hemopexin (HPX), haptoglobin, α1 microglobulin and albumin, with HPX having the highest affinity to CFH among these(3, 16).

We have reported the presence of elevated cell-free heme, in the plasma, lungs and alveolar fluid of mice and humans post exposure to the halogens, such as Cl_2_ and Bromine (Br_2_) (3, 5, 6, 8) as well as phosgene (4). However, detailed proteomics studies identifying key cellular processes and biological mechanisms responsible for the development of lung injury following exposure to Cl_2_ and their modification by hemopexin are lacking. Herein, we present new data showing that a single injection of human hemopexin in male and female adult C57BL/6 mice post Cl_2_ exposure decreases acute lung injury, mitochondrial injury and mortality. In addition, we used robust global-discovery proteomics in lung tissues to identify specific targets of Cl_2_ and its intermediates and the extent to which they are modified by human hemopexin injection and validated these findings with physiological and molecular biological techniques. Finally, In consideration of supply issues for human hemopexin, which is isolated from human plasma that hindered its promising clinical uses as a therapeutic agent for a number of diseases, including sickle cell disease, Acute Respiratory Distress Syndrome, trauma and hemorrhage (8, 19, 21, 29, 41, 47), we generated recombinant forms of human hemopexin in plants and assessed their pharmacokinetics efficacy in reversing acute lung injury in Cl_2_ exposed mice.

## Materials and Methods

### Animals

C57BL/6 8-12 weeks old male and female mice (20-25 g, body weight) mice were purchased from Charles River Laboratories (Wilmington, MA). All experimental procedures involving animals were approved by the University of Alabama at Birmingham (UAB) Institutional Animal Care and Use Committee (APN20990). Mice were allowed to acclimatize for at least three days prior to any experimental procedures. Those with elevated body temperatures, exhibiting hunched postures, refusing to eat and drink or with skin lesions were not used in this study

### Exposure to Cl_2_

Unanesthetized male and female mice were exposed to Cl_2_ (400 or 500 ppm for 30 min) in Plexiglas chambers between 7-10 AM as described previously (6, 26). According to UAB IACUC guidelines, all mice were injected with long acting buprenorphine (1 mg/kg subcutaneously) prior to exposure to minimize anxiety and discomfort. At the end of the exposure, the mice were returned to their cages and observed during the day at least every two h and once during the night. They were allowed access to food and water *ad libitum.* For the survival studies, mice were monitored for two weeks to determine the level of mortality between each group. Mice were sacrificed if their body weight dropped below 30% of their initial value.

### Recombinant Hemopexin

Recombinant hemopexin (prhHPX) was generated by AntoXa (Toronto, Canada).. A synthetic DNA sequence, corresponding to hHPX protein sequence, was cloned into an acceptor T-DNA vector downstream of a strong constitutive promoter. Vectors were transformed into *A. tumefaciens* EHA105 and glycerol seed stocks prepared. Greenhouse-grown 4-week old *N. benthamiana* KDFX plants were exposed to the prhHPX-transformed strains along with a proprietary system for suppression of post-translational gene silencing. After a further incubation for 7 days, the plants were harvested and protein extracted. The Hemopexin protein binds to Nickel-loaded resins such as Ni-Sepharose and is displaced by Imidazole. The product recovered from Ni-Sepharose was shown, by SDS-PAGE, to contain a significant amount of low- and high-molecular mass contaminants. These contaminants were resolved from the intact monomeric protein using size-exclusion chromatography on a Suprfrc S-200 column (Cytiva, Georgia, USA).

Mice were injected intramuscularly with either hHPX or prhHPX (5-10 µg/g BW in 100 µl of sterile normal saline) or an equal volume of sterile saline at 1 h after the cessation of Cl_2_ exposure while breathing room air. All physiological measurements were conducted at 24 h pos exposure while the mice were breathing room air. For pharmacokinetic studies, mice were sacrificed at various intervals post injection of hemopexin and blood samples were obtained by intracardial puncture. Plasma levels of hHPX and prhHPX were measured using an ELISA kit (product no. OKIA00066, AVIVA Systems Biology, San Diego, CA)) as described previously (6, 8). Non-encapsulated heme and hemoglobin concentrations were measured by using the QuantiChrom heme assay kit (Product No. DIHM-250; BioAssay Systems, Hayward, CA), according to the manufacturer’s instructions.

### Discovery Proteomics

Sample preparation and discovery proteomics analysis were carried out as described previously(1, 32). After sacrifice, lungs were perfused via a catheter inserted in the right ventricle and lavaged with normal saline till they were free of blood. Relative quantification across experiments were then performed via spectral counting, and when relevant, spectral count abundances were then normalized between samples.

### Biological Variables

For assessment of acute lung injury, mice were sacrificed 24 h post exposure to Cl_2_ or air and lavaged as described previously(7, 8) Total protein concentrations in cell-free lavage were measured with the BCA Protein Assay Kit (#23225; Thermo Fisher Scientific, Rockford, IL). Cells were re-suspended in 100μl of phosphate buffered saline (PBS) and counted using a Neubauer hemocytometer. Cells were then placed on slides using a Cellspin (Tharmac, Drosselweg, Germany) and stained using a two-stain set consisting of Eosin Y and a solution of thiazine dyes (Quik-Stain; Siemens, Washington, DC). Differential counts (specifically macrophage, neutrophils, and lymphocytes) were then performed on slides via light microscopy.

### RBC lysis

RBCs were isolated from blood samples from the abdominal aorta and subjected to mechanical stress by mixing them with glass beads; the mixtures were shaken for 2 hours (6). The concentrations of heme and hemoglobin in the plasma were measured by ELISA using an antibody against human HPX as described above. RBCs were then lysed with Triton X and this measurement was repeated. RBC fragmentation was then calculated as the ratio of (heme+hemoglobin) prior to and following the lysis with Triton X.

### Measurement of arterial blood gases (ABG)

Mice were anesthetized using isoflurane (5% for induction, 1-2% for maintenance) using room air as vehicle with attention being paid to maintain equal level of anesthesia (∼55-65 BPM breath rate). After laparotomy, the abdominal aorta was visualized and bluntly cleaned of fat tissue. A 25G needle, attached to a 1 mL syringe and loaded with 0.1 mL Heparin was inserted into the aorta and blood was drawn at a steady slow rate to avoid hemolysis and ensuring that there were no air bubbles aspirated. Approximately 0.1 mL blood is injected into a measurement cassette of a HESKA Element POC ABG instrument (Heska Corporation; Loveland, Colorado) to perform blood gas analysis.

### Lung Mitochondria DNA concentrations

The concentration of mtDNA in lung tissues was measured by two different methods. In the first set of measurements, a 25 mg section of lung tissue from the left lower lobe was harvested and lung DNA was extracted by using QIAamp DNA mini kit, (cat# 51306, Qiagen, Germany). The 8.1kb mtDNA was amplified by PCR and quantitatively analyzed by real-time PCR using specific primers provided in the Mouse Real-Time PCR Mitochondrial DNA Damage Analysis Kit from Detroit R&D (cat. #DD2M; Detroit, MI). Mitochondrial damage in the 8.1kb mtDNA fragment was quantified by a standard curve prepared. In the second set of measurements, DNA was amplified using the following 11Kb mitochondrial DNA specific primers, designed by one of the authors (SD): Mice-Mito-11Kb (F): CTT TAC GAG CCG TAG CCC AA; Mice-Mito-11Kb (R): AGC GAA GAA TCG GGT CAA GG; Mice-Mito-Short-(F): GGC CCA TTA AAC TTG GGG GT; Mice-Mito-Short-(R): TTA AGG GGA ACG TAT GGG CG. Real-time was performed by taking long PCR product as a template, to quantify the intact Mitochondrial DNA concentration with a mitochondrial DNA specific PCR primer (120bp long, designed by SD).

### Western blots

Western blots for the assessment of total and phosphoElF2a were performed using lung tissues processed for Discovery proteomics analysis. Lung tissues were homogenized with 3000 µl of T-PER Tissue Protein Extraction Reagent (PIERCE #78510, Rockford, IL) with protease and phosphatase inhibitor cocktail (Thermo Scientific #78442, Rockford, IL), and centrifuged at 10000xg for 5 minutes to pellet cell/tissue debris. The supernatant was collected and the protein concentration measured with the BCA Protein Assay (PIERCE #23228, Rockford, IL).

Subsequently, we loaded 25 µg protein of each sample to SDS-PAGE gels and incubated the membranes with 1:1000 eIF2α antibody (Cell Signaling Technology #9722, Danvers, MA) or 1:1000 Phospho-eIF2α antibody (Cell Signaling Technology #9721, Danvers, MA) in 5% w/v BSA, 1xTBS, 0.1% Tween 20 at 4°C with gentle shaking, overnight. Each membrane was then incubated with a β-Actin antibody (Santa Cruz Biotechnology #sc-47778, Dallas, Texas). Protein bands were quantified with ImageJ Lab Software.

### Statistical Analysis

Figures were generated and statistics performed using GraphPad Prism version 8 for Windows (GraphPad Software, San Diego, CA). The mean ± SEM was calculated in all experiments, and statistical significance was determined by either the one-way or the two-way ANOVA. In either case the Dunn–Šidák test for multiple comparisons was used for biological variable and the Bonferroni test for multiple comparisons of proteomic variables. Overall survival was analyzed by the Kaplan–Meier method. Differences in survival were tested for statistical significance by the log-rank test. A value of p < 0.05 was considered significant.

## Results

### Pharmacokinetic studies of hHPX

In the first set of experiments, we exposed adult male and female C57BL/6 mice to Cl_2_ (500 ppm for 30 m), returned them to room air and at 1h post exposure injected the mice with a bolus of human hemopexin (hHPX; 5 µg/ g BW) (**Figure 1A**). These data were fitted in a non-compartmental PK model (37). There were marked differences in the maximum concentration of hHPX in the plasma among air and Cl_2_ exposed mice (Cmax 9080±1482 vs. 2048±316 ng/ml; X± 1 SD; n=3; p<0.0013; unpaired two-tailed t-test) as well as in the daily area under the plasma drug concentration-time curve (AUC24; 174.6±21 vs. 49.6 at 24±5.3 hr*mg/L; X± 1 SD; n=3; p<0.0004; unpaired two-tailed t-test), which reflects the actual body exposure to hHPX. These differences are most likely due to the formation of heme-hHPX complexes and the elimination in the liver.

**Figure 1.**
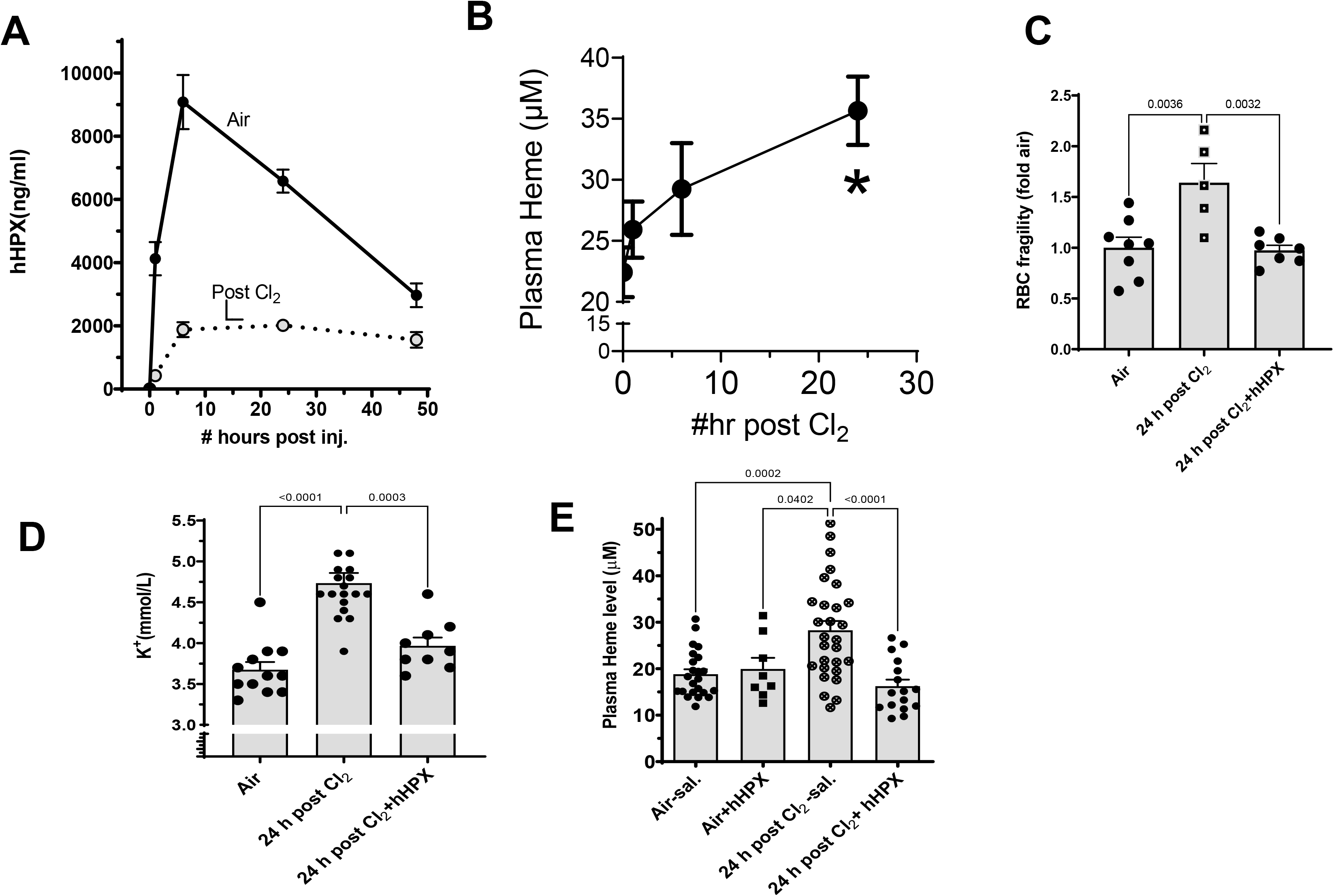
Injection of hHPX decreased heme levels and RBC fragility post Cl_2_. (A) C57BL/6 mice were exposed to air (solid circles) or Cl_2_ (500 ppm for 30 min; open circles connected by dotted line). One h post exposure they were injected intramuscularly with human hemopexin (hHPX; 5 µg/g BW). Mice were then sacrificed at the indicated times and the concentration of hHPX in their plasmas was measured by ELISA as described in the methods; n=3-5 for each condition. Corresponding mean values at each time point are significantly different from each other with the unpaired t-test (p<0.001). (**B)**. C57BL/6 mice were exposed to air or Cl_2_ (500 ppm for 30 min) and returned to room air. Measurements of plasma heme at 1, 6 or 24 h post exposure. Values are means ± 1 SEM; n=6-8 for each time point. * Significantly different from the air value (p<0.01) by Anova followed by the Dunn–Šidák test for multiple comparisons. The slope of the regression line is statistically different from zero (F=1.29; p=0.0022). (**C, D, E**) C57BL/6 mice were exposed to air or Cl_2_ (500 ppm for 30 min) and returned to room air. They were injected intramuscularly with human hemopexin (hHPX; 5 µg/g BW) at 1 h post exposure. Mice were then sacrificed at 24 h post exposure and the RBC fragility (**C)**, plasma K^+^ concentration (**D**) and plasma heme were measured as described in the methods. Each point represents a different animal. Means ± 1 SEM; statistical analysis by one way analysis of variance followed by the Dunn–Šidák test for multiple comparisons (GraphPad Prism 9).

### Injection of hHPX decreases heme levels

As shown in **Figure 1B**, plasma heme levels increased post exposure to Cl_2_; at 24 h mean concentration of plasma heme increased by about 60% as compared to the corresponding air value This is consistent with the in vitro measurement of RBC fragility (**Figure 1C**) and the large increase of plasma K^+^ seen 24 h post Cl_2_ (**Figure 1D**). A single injection of hHPX (5 µg/g BW) at 1 h post exposure, prevented RBC fractionation and decreased plasma heme and K^+^ concentrations to their corresponding air control values (**Figures 1C-E**). Injection of hHPX in air breathing mice had no effect on plasma heme (**Figure 1E**).

### Injection of hHPX decreases lung injury and prolongs survival

Exposure of C57BL/6 mice to Cl_2_ (500 ppm for 30 min) resulted in significant injury to the blood gas barrier, as shown by the large increase of BALF proteins (**Figure 2A**) and accumulation of inflammatory cells (in the BALF (**Figure 2B**); The fraction of neutrophils among BALF cells increased from 4±0.5 in air to 16.2±2 ×10^4^/ml at 24 h post Cl_2_ (X±1 SEM; n=4; p<0.001). At 24 h post Cl2 mice developed compensated respiratory acidosis as shown by the presence of severe hypercapnia (**Figure 2C**) and only a small change in pH from 7.35 in air to 7.29. At the same time, there was significant injury (to the mitochondrial DNA as shown by almost 40% decrease of intact mitochondrial DNA (**Figure 2D**) and an increase of lung mitochondrial DNA lesion (data not shown). A single injection of hHPX (5 µg/ g BW) at 1 h post exposure had a beneficial effect on all of these variables (**Figures 2A-2D**). As shown in **Figure 2E**, hHPX (which has a molecular weight of ∼ 60 kDa, similar to that of albumin) was one of the proteins detected in the BALF of mice at 24 h post Cl_2_ exposure.

**Figure 2.**
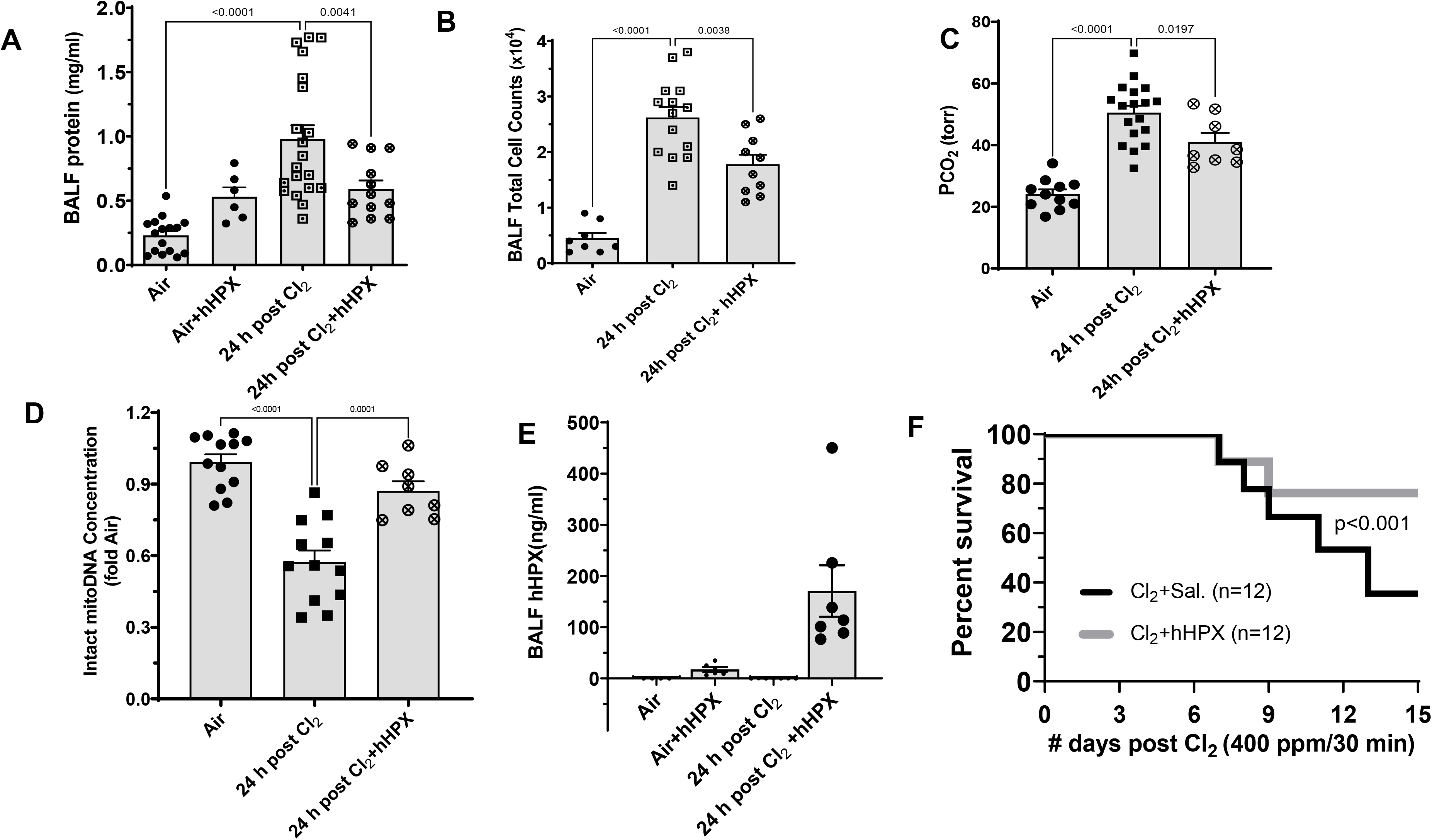
**Injection of hHPX decreased Cl_2_-induced acute lung injury and mortality**. (**A, B**). C57BL/6 mice were exposed to air or Cl_2_ (500 ppm for 30 min) and returned to room air. One h post exposure they were injected intramuscularly with human hemopexin (hHPX; 5 µg/g BW). Mice were then sacrificed at 24 h post exposure, their lungs were lavaged and the concentration of protein (**A)** and # of cells (**B**) in the BALF were measured as described in the text. Each point represents a different animal. Means ± 1 SEM; (**C**). C57BL/6 mice were exposed to air or Cl_2_ (500 ppm for 30 min) and returned to room air. One h post exposure they were injected intramuscularly with human hemopexin (hHPX; 5 µg/g BW). Their lungs were removed, and the concentration of mtDNA was measured by RT-PCR using Mitochondrial DNA Damage Analysis Kit from Detroit R&D (cat. #DD2M). Similar results were obtained when DNA was amplified using the following 11Kb mitochondrial DNA specific primers, designed by one of the authors (see METHODS). Means ± 1 SEM; statistical analysis by one way analysis of variance followed by the Dunn–Šidák for multiple comparisons. (**D**). hHPX in the BALF, measured by ELISA. Means ± 1 SEM; statistical analysis by one way analysis of variance followed by the Dunn–Šidák for multiple comparisons. (**E**) C57BL/6 mice were exposed to air or Cl_2_ (400 ppm for 30 min) and returned to room air. One h post exposure they were injected intramuscularly with human hemopexin (hHPX; 5 µg/g BW). Kaplan-Meir curves and statistical significance among them was calculated by GraphPad PRISM 9.1software.

In the next series of experiments, we exposed male and female C57BL/6 mice to 400 ppm Cl_2_ for 30 min which resulted in 50% mortality by 12 days (**Figure 2F**). A single injection of hHPX (5 µg/ g BW) at 1 h post exposure improved survival considerably: at 15 days post exposure less than 40% of mice injected with saline were alive vs. 80% of those receiving hHPX.

### Exposure to Cl_2_ induced large changes in the lung proteome

The unbiased global proteomics analysis of lung tissues showed that that exposure of mice to Cl_2_ resulted at 24 h later in the modification of 237 lung proteins: 92 of them were upregulated and 145 were downregulated (p<0.05, SAM>0.6, and Fold change>1.5). The top 80 proteins that were either increased or decreased two-fold at 24 h post Cl_2_ as compared to their corresponding air values and passed a 2-tiered statistical test are shown in **Tables 1A** and **1B.** The Heat Map and Principal Component Analysis, shown in **Figures 3 A-B** demonstrate clear changes among the air and 24 h post Cl_2_ lung proteomes. Furthermore, the pie chart in **Figure 3C** highlights the biological processes altered in the lungs of mice at 24 h post Cl_2_. The most prominent changes were the significant downregulations of the eukaryotic initiation factor 2 (elF2) which integrates a diverse array of stress-related signals to regulate both global and specific mRNA translation (48), coagulation and microRNA biogenesis signaling as well remodeling of epithelial adherence junctions.

**Table 1A.**
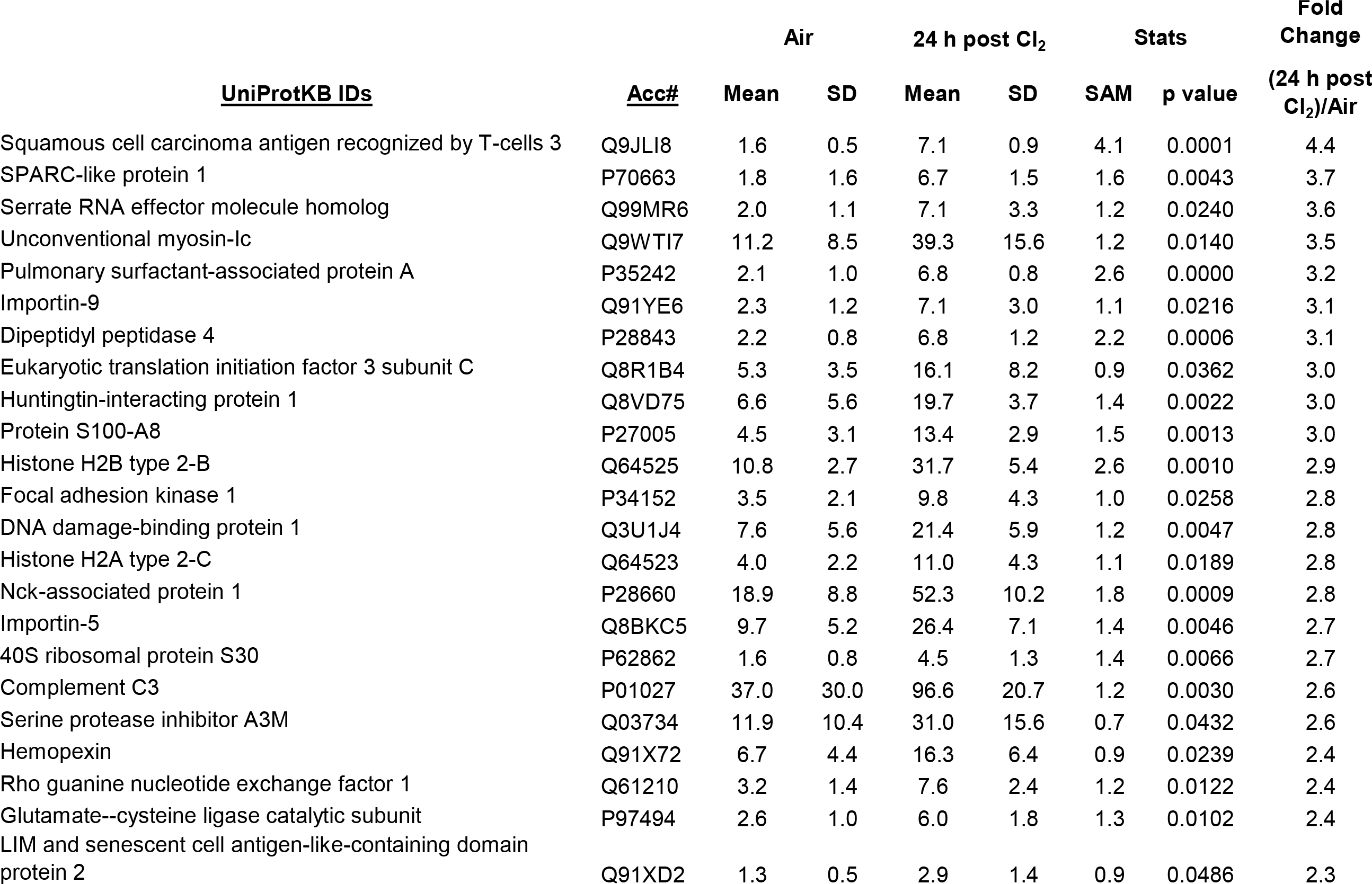
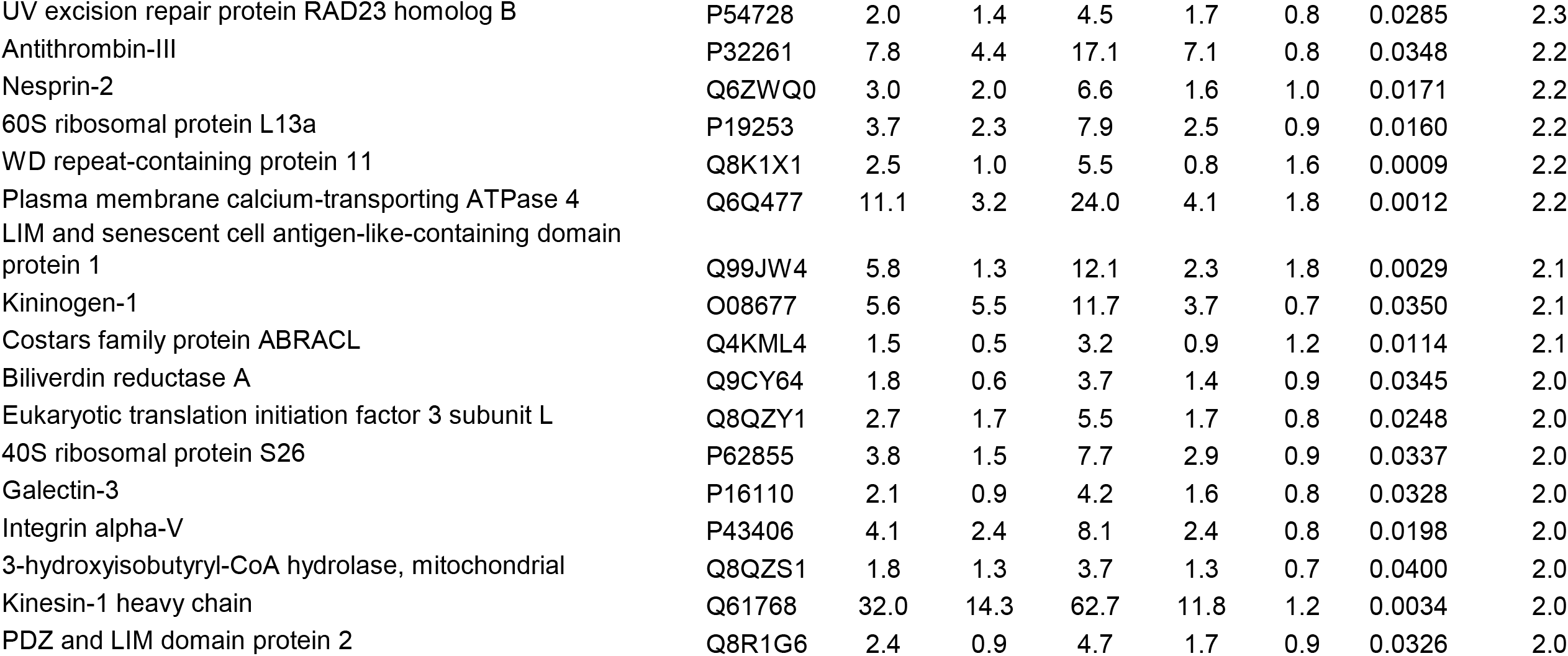
Top most increased lung proteins at 24 h post Cl_2._ All proteins have been GO annotated using a combination of the UniProtKB database, DAVID GO, and Genego Metacore. Spectral counts; Values are X ± 1 SD; n=6 for air and n=4 for 24 h post Cl_2_ p values obtained from non-parametric t-test analysis

| <u>UniProtKB IDs</u> | <u>Acc#</u> | <u>Air</u> |  | <u>24 h post Cl<sub>2</sub></u> |  | <u>Stats</u> |  | <u>Fold Change</u> |
| --- | --- | --- | --- | --- | --- | --- | --- | --- |
|  |  | <u>Mean</u> | <u>SD</u> | <u>Mean</u> | <u>SD</u> | <u>SAM</u> | <u>p value</u> | <u>(24 h post Cl<sub>2</sub>)/Air</u> |
| Squamous cell carcinoma antigen recognized by T-cells 3 | Q9JLI8 | 1.6 | 0.5 | 7.1 | 0.9 | 4.1 | 0.0001 | 4.4 |
| SPARC-like protein 1 | P70663 | 1.8 | 1.6 | 6.7 | 1.5 | 1.6 | 0.0043 | 3.7 |
| Serrate RNA effector molecule homolog | Q99MR6 | 2.0 | 1.1 | 7.1 | 3.3 | 1.2 | 0.0240 | 3.6 |
| Unconventional myosin-Ic | Q9WTI7 | 11.2 | 8.5 | 39.3 | 15.6 | 1.2 | 0.0140 | 3.5 |
| Pulmonary surfactant-associated protein A | P35242 | 2.1 | 1.0 | 6.8 | 0.8 | 2.6 | 0.0000 | 3.2 |
| Importin-9 | Q91YE6 | 2.3 | 1.2 | 7.1 | 3.0 | 1.1 | 0.0216 | 3.1 |
| Dipeptidyl peptidase 4 | P28843 | 2.2 | 0.8 | 6.8 | 1.2 | 2.2 | 0.0006 | 3.1 |
| Eukaryotic translation initiation factor 3 subunit C | Q8R1B4 | 5.3 | 3.5 | 16.1 | 8.2 | 0.9 | 0.0362 | 3.0 |
| Huntingtin-interacting protein 1 | Q8VD75 | 6.6 | 5.6 | 19.7 | 3.7 | 1.4 | 0.0022 | 3.0 |
| Protein S100-A8 | P27005 | 4.5 | 3.1 | 13.4 | 2.9 | 1.5 | 0.0013 | 3.0 |
| Histone H2B type 2-B | Q64525 | 10.8 | 2.7 | 31.7 | 5.4 | 2.6 | 0.0010 | 2.9 |
| Focal adhesion kinase 1 | P34152 | 3.5 | 2.1 | 9.8 | 4.3 | 1.0 | 0.0258 | 2.8 |
| DNA damage-binding protein 1 | Q3U1J4 | 7.6 | 5.6 | 21.4 | 5.9 | 1.2 | 0.0047 | 2.8 |
| Histone H2A type 2-C | Q64523 | 4.0 | 2.2 | 11.0 | 4.3 | 1.1 | 0.0189 | 2.8 |
| Nck-associated protein 1 | P28660 | 18.9 | 8.8 | 52.3 | 10.2 | 1.8 | 0.0009 | 2.8 |
| Importin-5 | Q8BKC5 | 9.7 | 5.2 | 26.4 | 7.1 | 1.4 | 0.0046 | 2.7 |
| 40S ribosomal protein S30 | P62862 | 1.6 | 0.8 | 4.5 | 1.3 | 1.4 | 0.0066 | 2.7 |
| Complement C3 | P01027 | 37.0 | 30.0 | 96.6 | 20.7 | 1.2 | 0.0030 | 2.6 |
| Serine protease inhibitor A3M | Q03734 | 11.9 | 10.4 | 31.0 | 15.6 | 0.7 | 0.0432 | 2.6 |
| Hemopexin | Q91X72 | 6.7 | 4.4 | 16.3 | 6.4 | 0.9 | 0.0239 | 2.4 |
| Rho guanine nucleotide exchange factor 1 | Q61210 | 3.2 | 1.4 | 7.6 | 2.4 | 1.2 | 0.0122 | 2.4 |
| Glutamate--cysteine ligase catalytic subunit | P97494 | 2.6 | 1.0 | 6.0 | 1.8 | 1.3 | 0.0102 | 2.4 |
| LIM and senescent cell antigen-like-containing domain protein 2 | Q91XD2 | 1.3 | 0.5 | 2.9 | 1.4 | 0.9 | 0.0486 | 2.3 |
| UV excision repair protein RAD23 homolog B | P54728 | 2.0 | 1.4 | 4.5 | 1.7 | 0.8 | 0.0285 | 2.3 |
| Antithrombin-III | P32261 | 7.8 | 4.4 | 17.1 | 7.1 | 0.8 | 0.0348 | 2.2 |
| Nesprin-2 | Q6ZWQ0 | 3.0 | 2.0 | 6.6 | 1.6 | 1.0 | 0.0171 | 2.2 |
| 60S ribosomal protein L13a | P19253 | 3.7 | 2.3 | 7.9 | 2.5 | 0.9 | 0.0160 | 2.2 |
| WD repeat-containing protein 11 | Q8K1X1 | 2.5 | 1.0 | 5.5 | 0.8 | 1.6 | 0.0009 | 2.2 |
| Plasma membrane calcium-transporting ATPase 4 | Q6Q477 | 11.1 | 3.2 | 24.0 | 4.1 | 1.8 | 0.0012 | 2.2 |
| LIM and senescent cell antigen-like-containing domain protein 1 | Q99JW4 | 5.8 | 1.3 | 12.1 | 2.3 | 1.8 | 0.0029 | 2.1 |
| Kininogen-1 | O08677 | 5.6 | 5.5 | 11.7 | 3.7 | 0.7 | 0.0350 | 2.1 |
| Costars family protein ABRACL | Q4KML4 | 1.5 | 0.5 | 3.2 | 0.9 | 1.2 | 0.0114 | 2.1 |
| Biliverdin reductase A | Q9CY64 | 1.8 | 0.6 | 3.7 | 1.4 | 0.9 | 0.0345 | 2.0 |
| Eukaryotic translation initiation factor 3 subunit L | Q8QZY1 | 2.7 | 1.7 | 5.5 | 1.7 | 0.8 | 0.0248 | 2.0 |
| 40S ribosomal protein S26 | P62855 | 3.8 | 1.5 | 7.7 | 2.9 | 0.9 | 0.0337 | 2.0 |
| Galectin-3 | P16110 | 2.1 | 0.9 | 4.2 | 1.6 | 0.8 | 0.0328 | 2.0 |
| Integrin alpha-V | P43406 | 4.1 | 2.4 | 8.1 | 2.4 | 0.8 | 0.0198 | 2.0 |
| 3-hydroxyisobutyryl-CoA hydrolase, mitochondrial | Q8QZS1 | 1.8 | 1.3 | 3.7 | 1.3 | 0.7 | 0.0400 | 2.0 |
| Kinesin-1 heavy chain | Q61768 | 32.0 | 14.3 | 62.7 | 11.8 | 1.2 | 0.0034 | 2.0 |
| PDZ and LIM domain protein 2 | Q8R1G6 | 2.4 | 0.9 | 4.7 | 1.7 | 0.9 | 0.0326 | 2.0 |

**Table 1B.** Top most **decreased** lung proteins at 24 h post C_l2_. All proteins have been GO annotated using a combination of the UniProtKB database, DAVID GO, and Genego Metacore.

| UniProtKB IDs | Acc# | Air |  | 24 h post Cl<sub>2</sub> |  | Stats |  | Fold change<br>24 h post Cl<sub>2</sub>/Air |
| --- | --- | --- | --- | --- | --- | --- | --- | --- |
|  |  | Mean | SD | Mean | SD | SAM | t-test |  |
| V-type proton ATPase subunit E 1 | P50518 | 6.8 | 3.1 | 2.9 | 2.1 | 0.7 | 0.0227 | -2.4 |
| Valacyclovir hydrolase | Q8R164 | 8.2 | 2.8 | 3.4 | 1.1 | 1.2 | 0.0035 | -2.4 |
| 60S ribosomal protein L21 | O09167 | 3.2 | 1.8 | 1.3 | 0.6 | 0.8 | 0.0270 | -2.4 |
| Lipoamide acyltransferase component of branched-chain alpha-keto acid dehydrogenase complex, mitochondrial | P53395 | 4.4 | 1.6 | 1.8 | 1.0 | 1.0 | 0.0065 | -2.4 |
| Carbonyl reductase [NADPH] 3 | Q8K354 | 5.8 | 2.2 | 2.4 | 1.1 | 1.0 | 0.0060 | -2.4 |
| Protein dopey-2 | Q3UHQ6 | 3.9 | 0.9 | 1.6 | 0.6 | 1.5 | 0.0032 | -2.4 |
| Heterogeneous nuclear ribonucleoprotein U | Q8VEK3 | 34.4 | 7.3 | 13.8 | 6.8 | 1.5 | 0.0013 | -2.5 |
| Structural maintenance of chromosomes protein 1A | Q9CU62 | 7.7 | 3.5 | 3.1 | 1.0 | 1.0 | 0.0115 | -2.5 |
| Astrocytic phosphoprotein PEA-15 | Q62048 | 8.6 | 3.5 | 3.4 | 1.6 | 1.0 | 0.0076 | -2.5 |
| 40S ribosomal protein S23 | P62267 | 12.6 | 4.5 | 5.0 | 1.7 | 1.2 | 0.0038 | -2.5 |
| Dipeptidyl peptidase 1 | P97821 | 2.7 | 1.9 | 1.1 | 0.0 | 0.9 | 0.0420 | -2.6 |
| Short/branched chain specific acyl-CoA dehydrogenase, mitochondrial | Q9DBL1 | 5.5 | 2.5 | 2.1 | 0.1 | 1.3 | 0.0101 | -2.6 |
| Heterogeneous nuclear ribonucleoprotein Q | Q7TMK9 | 4.7 | 0.9 | 1.8 | 1.3 | 1.3 | 0.0212 | -2.6 |
| L-lactate dehydrogenase B chain | P16125 | 25.2 | 16.1 | 9.5 | 0.4 | 1.0 | 0.0310 | -2.7 |
| Collagen alpha-1(XIV) chain | Q80X19 | 16.9 | 7.7 | 6.3 | 4.2 | 0.9 | 0.0115 | -2.7 |
| Heterogeneous nuclear ribonucleoprotein U-like protein 2 | Q00PI9 | 5.6 | 1.4 | 2.1 | 1.1 | 1.4 | 0.0035 | -2.7 |
| U5 small nuclear ribonucleoprotein 200 kDa helicase | Q6P4T2 | 15.7 | 5.5 | 5.8 | 3.2 | 1.1 | 0.0036 | -2.7 |
| Dolichyl-diphosphooligosaccharide--protein glycosyltransferase subunit 1 | Q91YQ5 | 3.6 | 1.3 | 1.3 | 0.6 | 1.2 | 0.0027 | -2.7 |
| Isoleucine--tRNA ligase, mitochondrial | Q8BIJ6 | 6.5 | 1.4 | 2.4 | 1.0 | 1.7 | 0.0003 | -2.8 |
| Poly(rC)-binding protein 2 | Q61990 | 22.1 | 8.4 | 7.9 | 3.8 | 1.2 | 0.0038 | -2.8 |
| Glutathione S-transferase theta-3 | Q99L20 | 13.3 | 4.1 | 4.7 | 1.2 | 1.6 | 0.0012 | -2.8 |
| Nucleobindin-1 | Q02819 | 4.0 | 2.3 | 1.4 | 0.6 | 0.9 | 0.0195 | -2.9 |
| TAR DNA-binding protein 43 | Q921F2 | 9.4 | 5.2 | 3.2 | 1.3 | 1.0 | 0.0151 | -3.0 |
| Chromodomain-helicase-DNA-binding protein 4 | Q6PDQ2 | 10.3 | 2.1 | 3.4 | 1.7 | 1.8 | 0.0003 | -3.1 |
| Heterogeneous nuclear ribonucleoprotein D0 | Q60668 | 19.1 | 4.8 | 5.7 | 1.8 | 2.0 | 0.0002 | -3.3 |
| Receptor expression-enhancing protein 5 | Q60870 | 9.8 | 4.9 | 2.9 | 1.7 | 1.0 | 0.0082 | -3.4 |
| Receptor-type tyrosine-protein phosphatase C | P06800 | 7.4 | 2.9 | 2.1 | 0.9 | 1.4 | 0.0025 | -3.5 |
| Interleukin enhancer-binding factor 2 | Q9CXY6 | 3.8 | 0.4 | 1.1 | 0.0 | 5.9 | 0.0000 | -3.6 |
| Enhancer of mRNA-decapping protein 4 | Q3UJB9 | 5.2 | 0.8 | 1.4 | 0.7 | 2.6 | 0.0006 | -3.6 |
| Splicing factor, proline- and glutamine-rich | Q8VIJ6 | 19.6 | 6.3 | 4.9 | 3.1 | 1.6 | 0.0007 | -4.0 |
| Lysozyme C-2 | P08905 | 42.6 | 16.3 | 10.5 | 1.6 | 1.8 | 0.0023 | -4.1 |
| Myb-binding protein 1A | Q7TPV4 | 16.2 | 4.3 | 3.9 | 2.5 | 1.8 | 0.0002 | -4.2 |
| Prostaglandin E synthase 3 | Q9R0Q7 | 9.4 | 6.1 | 2.1 | 1.0 | 1.0 | 0.0154 | -4.6 |
| Phospholipid hydroperoxide glutathione peroxidase | O70325 | 10.9 | 0.8 | 2.4 | 1.3 | 4.2 | 0.0001 | -4.6 |
| Heterogeneous nuclear ribonucleoprotein A3 | Q8BG05 | 25.0 | 6.0 | 5.2 | 2.0 | 2.5 | 0.0001 | -4.8 |
| Splicing factor 3B subunit 1 | Q99NB9 | 15.9 | 5.0 | 3.2 | 1.5 | 1.9 | 0.0005 | -5.0 |
| ATP-dependent RNA helicase A | O70133 | 43.7 | 8.3 | 8.3 | 2.8 | 3.2 | 0.0000 | -5.2 |
| Proteolipid protein 2 | Q9R1Q7 | 11.1 | 2.9 | 1.8 | 1.0 | 2.4 | 0.0001 | -6.1 |
| E3 ubiquitin-protein ligase RNF213 | E9Q555 | 9.0 | 3.5 | 1.4 | 0.6 | 1.9 | 0.0013 | -6.5 |
| Proteasome subunit beta type-9 | P28076 | 9.1 | 3.3 | 1.3 | 0.5 | 2.0 | 0.0010 | -7.0 |

**Figure 3.**
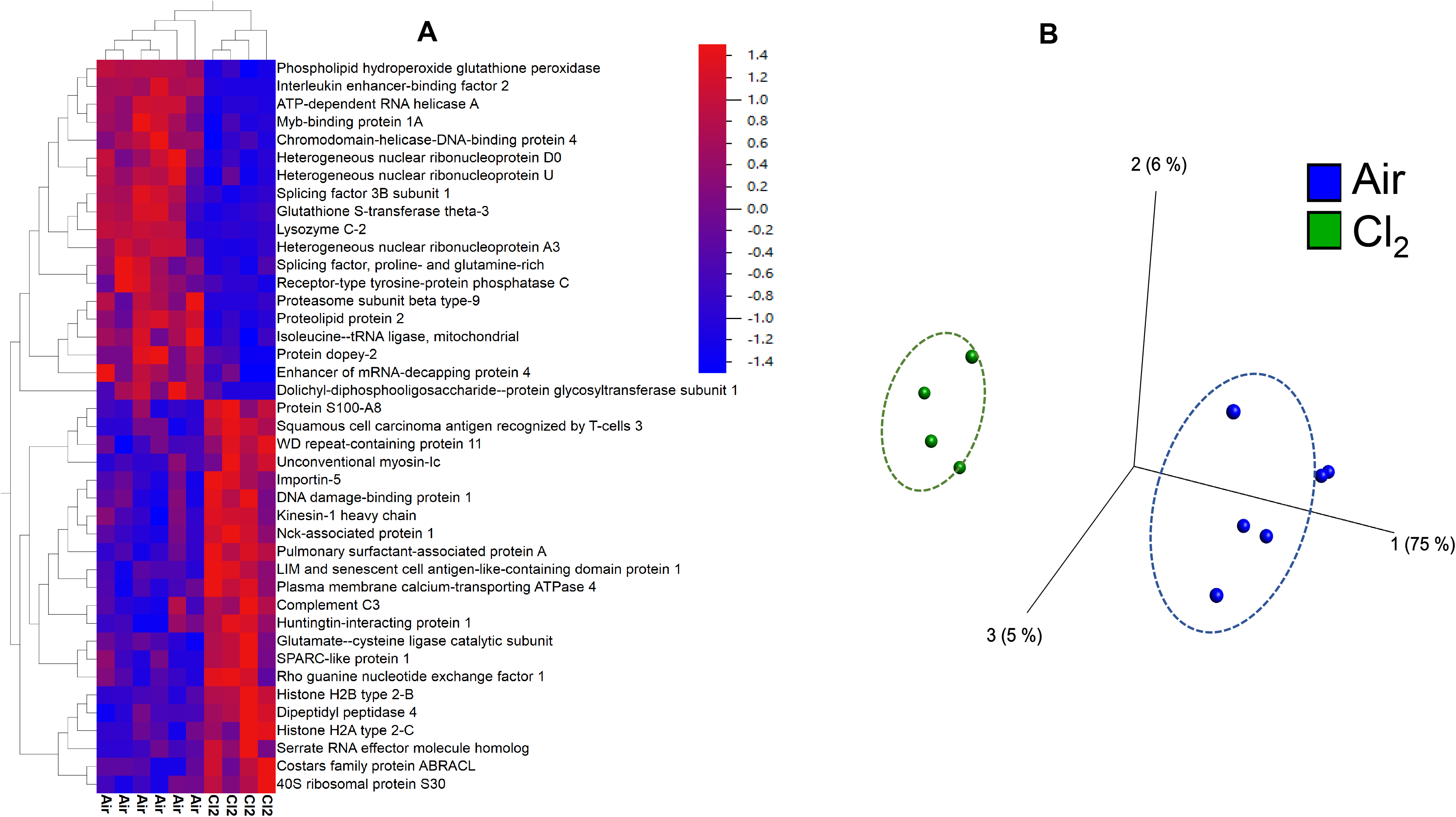
Heat Map and PCA Analysis of Lung Proteome post Cl_2_. C57BL/6 mice were exposed to air or Cl_2_ (500 ppm for 30 min) and returned to room air. Twenty four hours later the mice were sacrificed and their lungs were removed and proteins were processed for global proteomics analysis as discussed in the METHODS. The top 40 proteins (shown in Table 2) that passed a 2-tiered statistics test with p=0.01 and q=0.02 and were changed at least two fold as compared to air were used for these analyses. (A): the 2-dimensional hierarchical analysis heat map demonstrates which proteins are increased (red) or decreased (blue) in Cl_2_-vs. air-treated mice. Each column represents data from a different mouse (n=6air; n=4 24 h post Cl_2_). (B): PCA complements the heat map by using a similar cluster approach that determines which animal (based on protein quantification for all proteins in the top list) is similar across all animals analyzed. Notice tight clustering for air and Cl_2_-exposed mice (yellow and red) with a clear separation between the 2 groups. Each symbol indicates a different mouse. (n=6 air; n=4 24 h post Cl_2_).The numbers in parenthesis show the variations among each principal axis (C). Top process networks pie chart from systems biology analysis carried out on the most significantly changed lung proteins at 24 h post exposure of mice to Cl_2_.

### hHPX reverses Cl_2_-induced changes in the lung proteome

Eighty-three of the 237 lung proteins that were significantly changed (increased or decreased) in the lungs of mice at 24 h post Cl_2_ were modified by a single injection of hHPX at 1h post exposure (**Table 2**). The two dimensional (2D) hierarchical heat map (HCA-HM, **Figure 4A**), the principal component analysis (PCA, **Figure 4B**) of the three groups of data (air, Cl_2_, Cl_2_+hHPX), demonstrate clear differences in the lung proteomes of mice exposed to Cl_2_ and treated with hHPX vs. saline. Injection of hHPX in air breathing mice resulted in modification of significant number of proteins and physiological processes (**Supplemental Figures 1A,B**), in spite of the fact that hHPX did not alter plasma heme levels in air breathing mice.

**Table 2.**
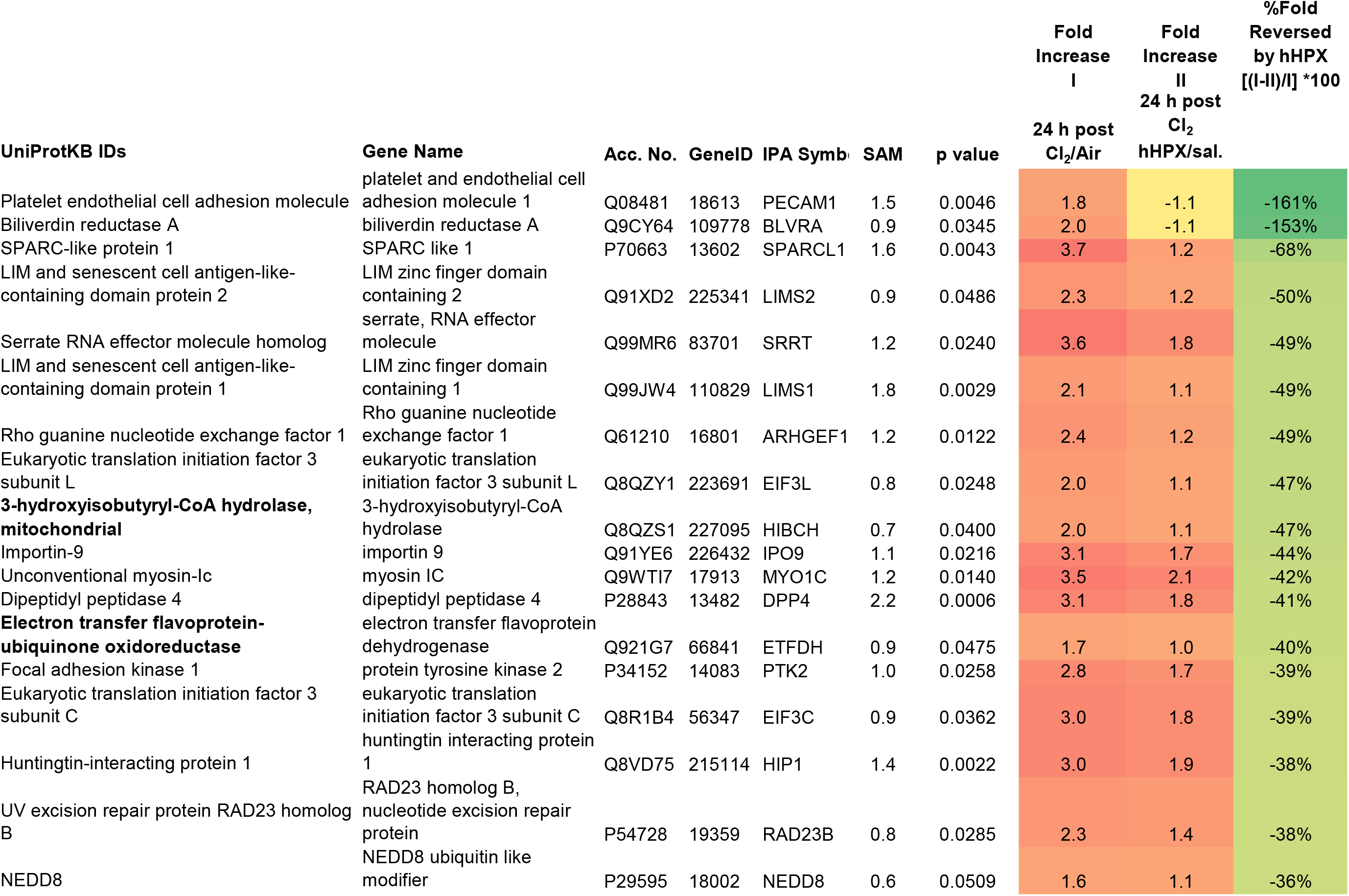
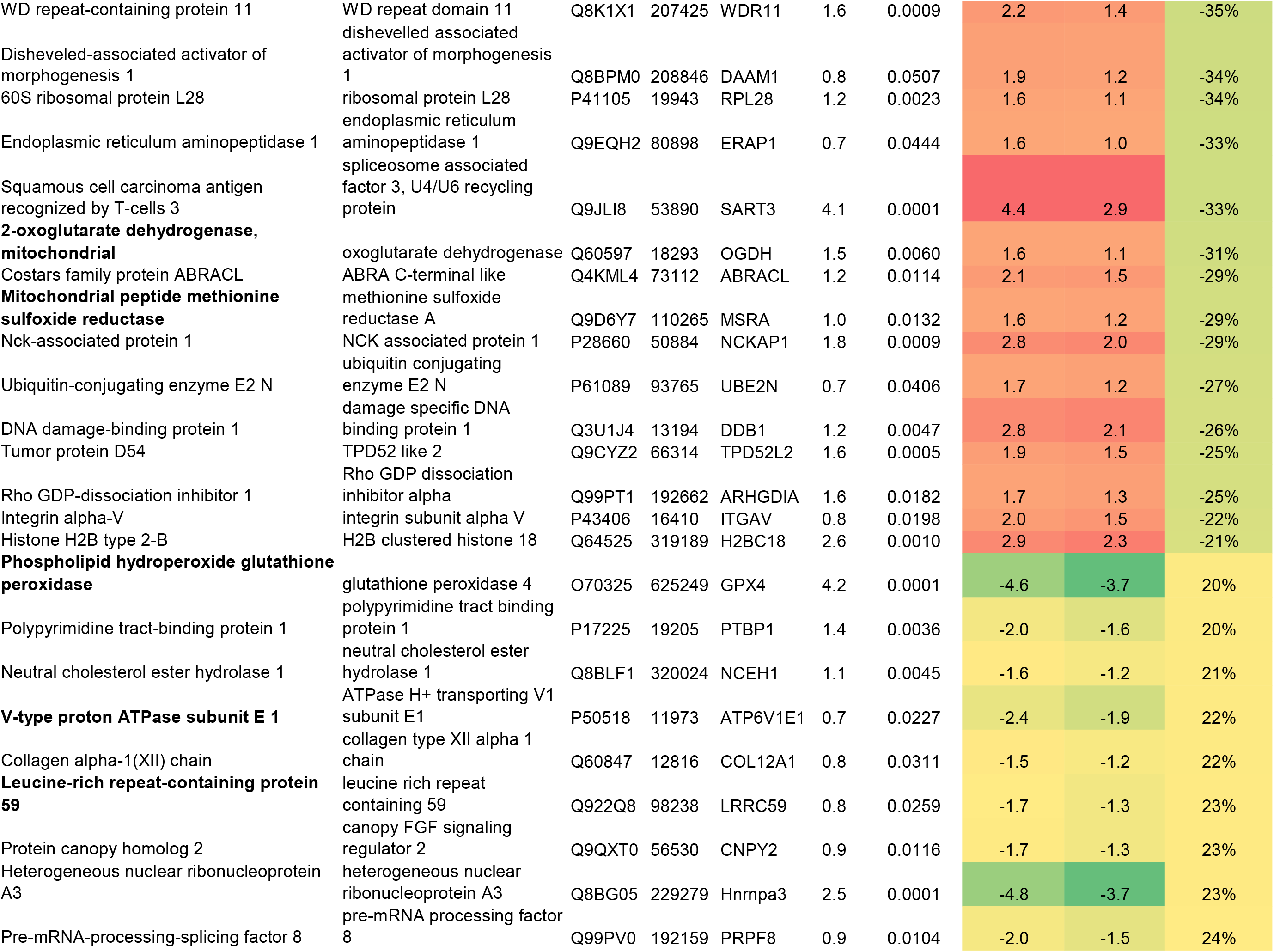
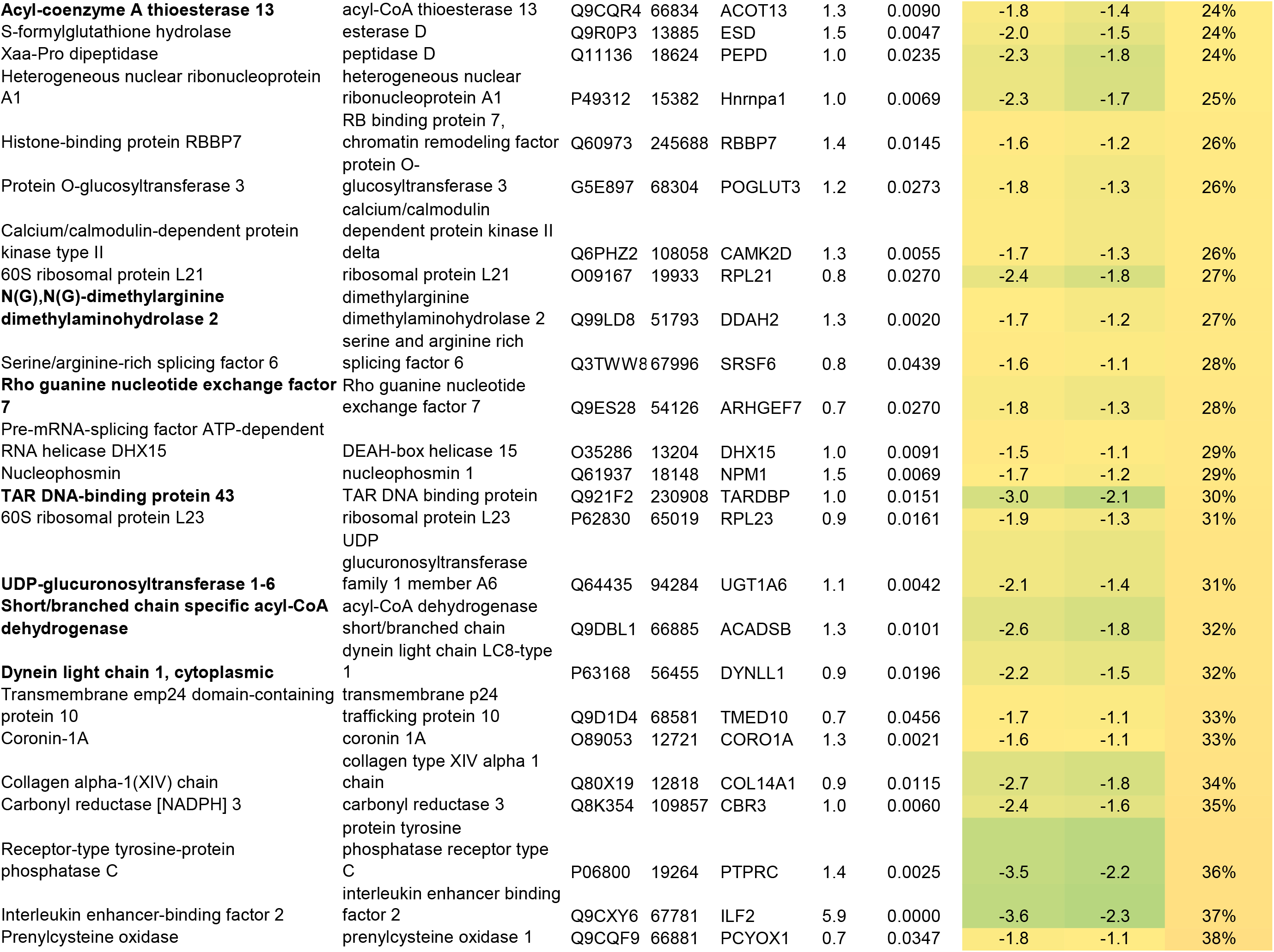
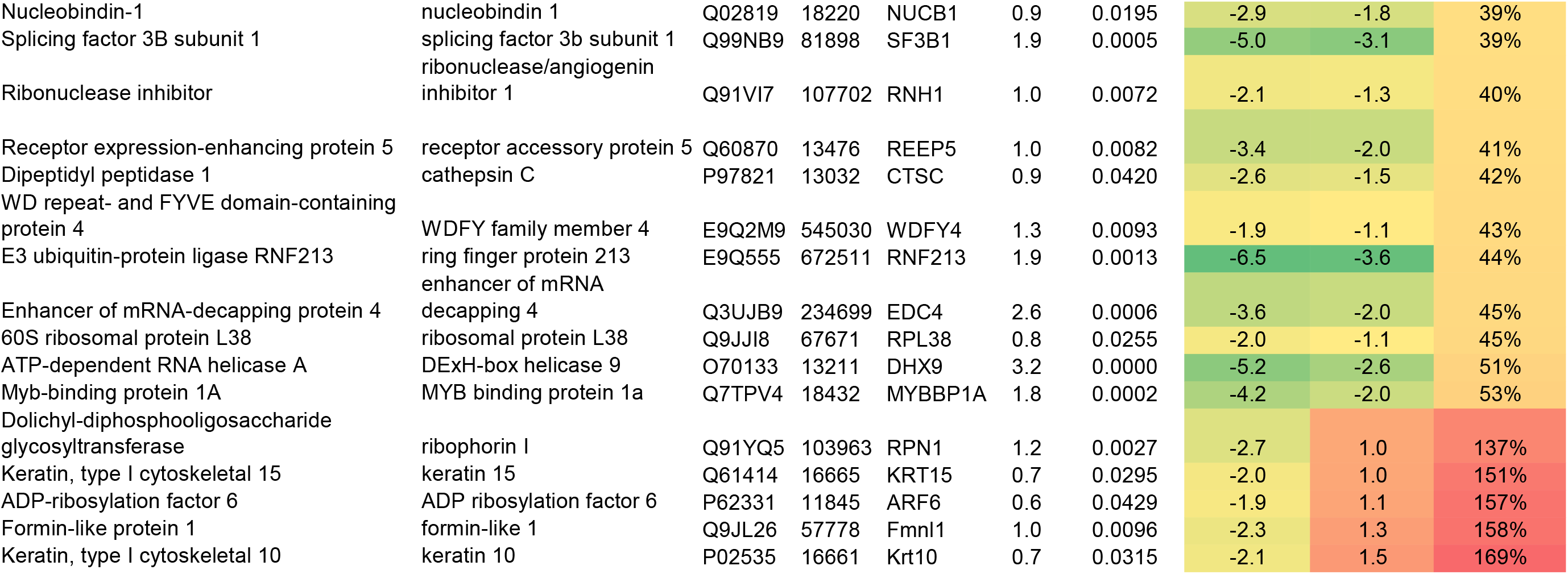
Mice were exposed to Cl_2_ (500 ppm for 30 min) and returned to room air. Twenty four h later, they were sacrificed and lung tissue was processed for LCMS2 analysis as described in the METHODS. The top 40 protein that were increased (highlighted in red) and decreased (green) following exposure to Cl_2_ are shown. Mean values ±1 SDV from four mice in each group.

**Figure 4.**
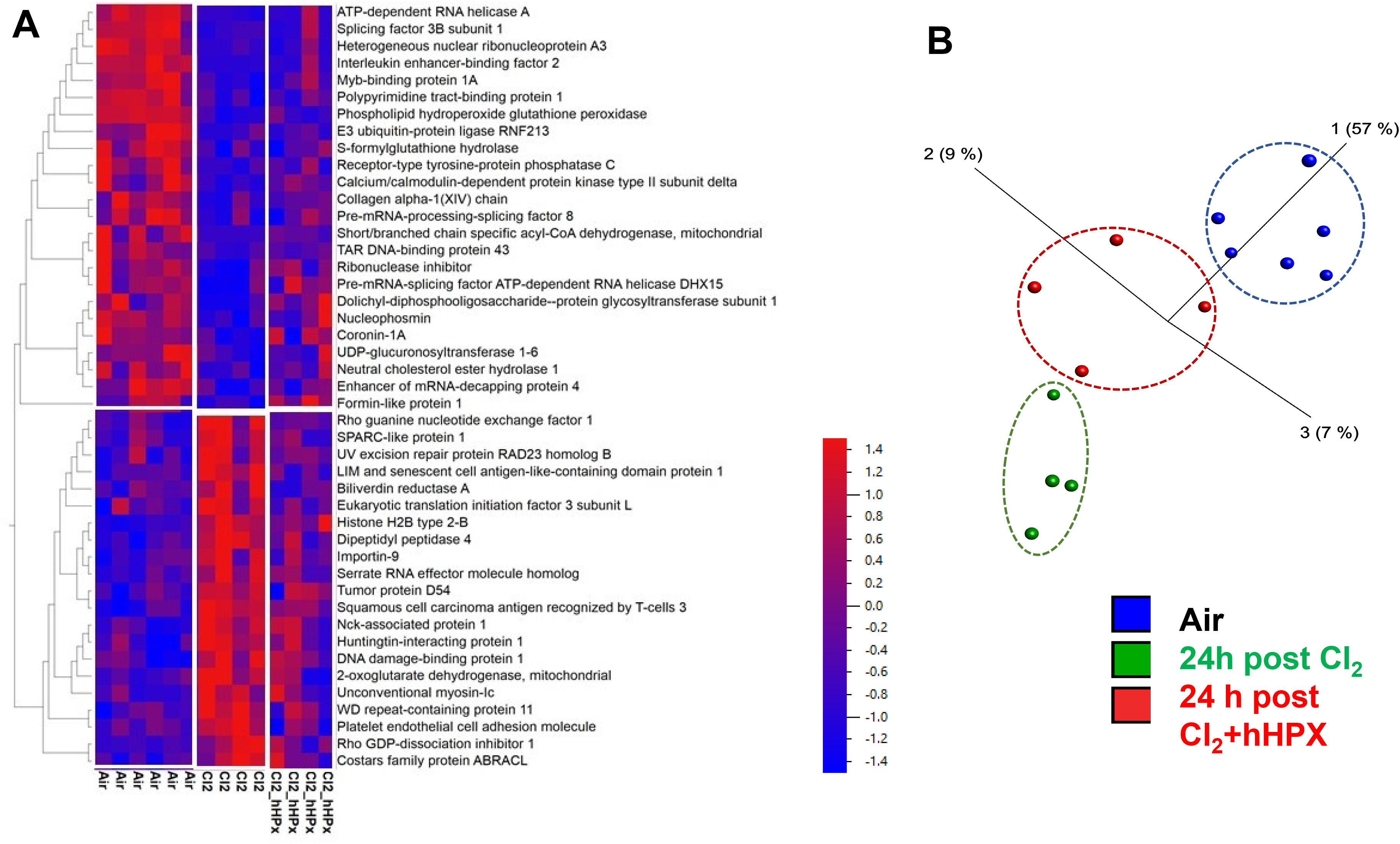
PCA Analysis and Heat Map post Cl_2_ and hHPX. C57BL/6 mice were exposed to air or Cl_2_ (500 ppm for 30 min) and returned to room air. One hour later they received a single dose of human hemopexin (5 µg/ g BW) intramuscularly. Twenty four hours later the mice were sacrificed and their lungs were removed and proteins were processed for global proteomics analysis as discussed in the METHODS. (**A**). The 2-dimensional hierarchical analysis heat map demonstrates which proteins are increased (red) or decreased (blue) in Cl2-vs. air and Cl_2_+hHPX vs Cl_2_ (q<0.1, p<0.05, top 46). (**B**) PCA complements the heat map by using a similar cluster approach that determines which animal (based on protein quantification for all proteins in the top list) is similar across all animals analyzed. Notice tight clustering for air, Cl_2_ and Cl_2_+hHPX mice with a clear separation between the three groups. Each point indicates a different mouse. (**C**). Volcano plot of the log10 P value vs. log2 fold change (Cl_2_ vs. Cl_2_+hHPX) demonstrating the distribution of the entire data set of proteins with upper limits (above the line) indicating statistically significant changes and outer limits (to the right and left of each line) indicating significant fold changes as outlined in materials and methods under statistics. Note that although fold change is visualized as log2, the cutoff value of ±1.5 was applied to the fold change before logging, thereby yielding the indicated ±0.6 limits.

### Biological Processes and Pathways via Systems Biology

To identify key biological processes altered following post treatment of Cl_2_ exposed mice with hHPX, we performed in-depth systems biology analysis using the data listed in **Table 2**. The top pathways modified by hHPX in the proteome of Cl_2_ exposed mice are shown in **Table 3** and in the two 2D bubble diagram in **Figure 5**. This analysis suggests that hHPX treatment modified a number of important processes associated with eIF2, Integrin signaling and mitochondrial dysfunction.

**Table 3.**

| **<u>Ingenuity Canonical Pathways</u>** | **<u>-log(p-value)</u>** | **<u>Ratio</u>** | **<u>Molecules</u>** |
| --- | --- | --- | --- |
| EIF2 Signaling | 4.93 | 0.0308 | EIF3C,EIF3L,PTBP1,RPL21,RPL23,RPL28,RPL38 |
| NER (Nucleotide Excision Repair, Enhanced Pathway) | 3.61 | 0.0444 | DDB1,NEDD8,RAD23B,UBE2N |
| Paxillin Signaling | 3.32 | 0.0374 | ARF6,ARHGEF7,ITGAV,PTK2 |
| Integrin Signaling | 3.12 | 0.0236 | ARF6,ARHGEF7,ITGAV,LIMS1,PTK2 |
| Actin Cytoskeleton Signaling | 2.85 | 0.0205 | ARHGEF1,ARHGEF7,ITGAV,NCKAP1,PTK2 |
| Valine Degradation I | 2.69 | 0.1 | ACADSB,HIBCH |
| RHOGDI Signaling | 2.18 | 0.0182 | ARHGDIA,ARHGEF1,ARHGEF7,ITGAV |
| Formaldehyde Oxidation II (Glutathione-dependent) | 2.17 | 0.5 | ESD |
| Thrombin Signaling | 2.15 | 0.0178 | ARHGEF1,ARHGEF7,CAMK2D,PTK2 |
| PAK Signaling | 2.14 | 0.0256 | ARHGEF7,ITGAV,PTK2 |
| Ferroptosis Signaling Pathway | 2 | 0.0227 | ARF6,DPP4,GPX4 |
| RAC Signaling | 1.95 | 0.0219 | ITGAV,NCKAP1,PTK2 |
| Reelin Signaling in Neurons | 1.94 | 0.0217 | ARHGEF1,ARHGEF7,CAMK2D |
| Spliceosomal Cycle | 1.92 | 0.0408 | DHX15,SF3B1 |
| Signaling by Rho Family GTPases | 1.9 | 0.015 | ARHGEF1,ARHGEF7,ITGAV,PTK2 |
| 2-ketoglutarate Dehydrogenase Complex | 1.87 | 0.25 | OGDH |
| Heme Degradation | 1.87 | 0.25 | BLVRA |
| CSDE1 Signaling Pathway | 1.81 | 0.0357 | EDC4,PTBP1 |
| Phagosome Maturation | 1.75 | 0.0184 | ATP6V1E1,CTSC,DYNLL1 |
| Molecular Mechanisms of Cancer | 1.74 | 0.0111 | ARHGEF1,ARHGEF7,CAMK2D,ITGAV,PTK2 |
| Agrin Interactions at Neuromuscular Junction | 1.64 | 0.029 | ARHGEF7,PTK2 |
| Regulation of eIF4 and p70S6K Signaling | 1.63 | 0.0166 | EIF3C,EIF3L,ITGAV |
| Pulmonary Fibrosis Idiopathic Signaling Pathway | 1.61 | 0.0123 | COL12A1,ITGAV,PTK2,RBBP7 |
| Glioma Invasiveness Signaling | 1.6 | 0.0274 | ITGAV,PTK2 |
| Xenobiotic Metabolism PXR Signaling Pathway | 1.56 | 0.0155 | CAMK2D,ESD,UGT1A6 |
| Mitochondrial Dysfunction | 1.54 | 0.0116 | CAMK2D,GPX4,OGDH,TARDBP |
| ILK Signaling | 1.52 | 0.0149 | LIMS1,LIMS2,PTK2 |
| Chemokine Signaling | 1.51 | 0.0247 | CAMK2D,PTK2 |
| Clathrin-mediated Endocytosis Signaling | 1.48 | 0.0144 | ARF6,HIP1,PCYOX1 |
| Regulation of Cellular Mechanics by Calpain Protease | 1.43 | 0.0222 | ITGAV,PTK2 |
| Isoleucine Degradation I | 1.25 | 0.0588 | ACADSB |
| Regulation of Actin-based Motility by Rho | 1.24 | 0.0174 | ARHGDIA,ITGAV |
| RHOA Signaling | 1.18 | 0.0161 | ARHGEF1,PTK2 |
| GP6 Signaling Pathway | 1.16 | 0.0157 | COL12A1,PTK2 |
| HGF Signaling | 1.14 | 0.0152 | ITGAV,PTK2 |
| Gα12/13 Signaling | 1.13 | 0.015 | ARHGEF1,PTK2 |
| TCA Cycle II (Eukaryotic) | 1.13 | 0.0435 | OGDH |
| Xenobiotic Metabolism Signaling | 1.12 | 0.0103 | CAMK2D,ESD,UGT1A6 |
| Glutathione Redox Reactions I | 1.04 | 0.0357 | GPX4 |
| PTEN Signaling | 1.04 | 0.0132 | ITGAV,PTK2 |
| Role of p14/p19ARF in Tumor Suppression | 1.02 | 0.0333 | NPM1 |
| Nucleotide Excision Repair Pathway | 0.953 | 0.0286 | RAD23B |
| Pyrimidine Ribonucleotides Interconversion | 0.909 | 0.0256 | DHX9 |
| Thyroid Hormone Metabolism II (via Conjugation and/or D | 0.888 | 0.0244 | UGT1A6 |
| Pyrimidine Ribonucleotides De Novo Biosynthesis | 0.879 | 0.0238 | DHX9 |
| GNRH Signaling | 0.867 | 0.0105 | CAMK2D,PTK2 |
| Leukocyte Extravasation Signaling | 0.86 | 0.0104 | PECAM1,PTK2 |
| PI3K/AKT Signaling | 0.835 | 0.01 | ITGAV,LIMS1 |
| Ephrin A Signaling | 0.833 | 0.0213 | PTK2 |
| Retinol Biosynthesis | 0.833 | 0.0213 | ESD |
| Ephrin Receptor Signaling | 0.828 | 0.0099 | ITGAV,PTK2 |
| Role of Tissue Factor in Cancer | 0.812 | 0.00966 | ITGAV,PTK2 |
| IL-8 Signaling | 0.802 | 0.00952 | ITGAV,PTK2 |
| Protein Kinase A Signaling | 0.792 | 0.0073 | CAMK2D,PTK2,PTPRC |
| mTOR Signaling | 0.789 | 0.00935 | EIF3C,EIF3L |
| ERK/MAPK Signaling | 0.786 | 0.0093 | ITGAV,PTK2 |
| Multiple Sclerosis Signaling Pathway | 0.764 | 0.00901 | RBBP7,RNF213 |
| Primary Immunodeficiency Signaling | 0.743 | 0.0169 | PTPRC |
| Nicotine Degradation III | 0.743 | 0.0169 | UGT1A6 |
| PCP (Planar Cell Polarity) Pathway | 0.736 | 0.0167 | DAAM1 |
| Semaphorin Signaling in Neurons | 0.73 | 0.0164 | PTK2 |
| Osteoarthritis Pathway | 0.724 | 0.00847 | ITGAV,MYBBP1A |
| Melatonin Degradation I | 0.717 | 0.0159 | UGT1A6 |
| Nicotine Degradation II | 0.693 | 0.0149 | UGT1A6 |
| Superpathway of Melatonin Degradation | 0.688 | 0.0147 | UGT1A6 |
| Remodeling of Epithelial Adherens Junctions | 0.688 | 0.0147 | ARF6 |
| GM-CSF Signaling | 0.676 | 0.0143 | CAMK2D |
| Melatonin Signaling | 0.666 | 0.0139 | CAMK2D |
| Ephrin B Signaling | 0.666 | 0.0139 | PTK2 |
| Serotonin Degradation | 0.666 | 0.0139 | UGT1A6 |
| Caveolar-mediated Endocytosis Signaling | 0.65 | 0.0133 | ITGAV |
| Angiopoietin Signaling | 0.645 | 0.0132 | PTK2 |
| Hypoxia Signaling in the Cardiovascular System | 0.645 | 0.0132 | UBE2N |
| Macropinocytosis Signaling | 0.645 | 0.0132 | ARF6 |
| NF-κB Activation by Viruses | 0.635 | 0.0128 | ITGAV |
| IL-7 Signaling Pathway | 0.635 | 0.0128 | PTK2 |
| Maturity Onset Diabetes of Young (MODY) Signaling | 0.63 | 0.0127 | OGDH |
| Axonal Guidance Signaling | 0.608 | 0.00588 | ARHGEF7,ITGAV,PTK2 |
| Xenobiotic Metabolism AHR Signaling Pathway | 0.594 | 0.0115 | UGT1A6 |
| Crosstalk between Dendritic Cells and Natural Killer Cells | 0.577 | 0.011 | CAMK2D |
| Apelin Adipocyte Signaling Pathway | 0.577 | 0.011 | GPX4 |
| Pyroptosis Signaling Pathway | 0.569 | 0.0108 | DHX9 |
| Actin Nucleation by ARP-WASP Complex | 0.569 | 0.0108 | ITGAV |
| Fcγ Receptor-mediated Phagocytosis in Macrophages and Neutrophils | 0.565 | 0.0106 | ARF6 |
| Cardiac Hypertrophy Signaling (Enhanced) | 0.559 | 0.00554 | CAMK2D,ITGAV,PTK2 |
| Small Cell Lung Cancer Signaling | 0.557 | 0.0104 | PTK2 |
| DNA Methylation and Transcriptional Repression Signaling | 0.55 | 0.0102 | RBBP7 |
| VEGF Signaling | 0.546 | 0.0101 | PTK2 |
| Synaptogenesis Signaling Pathway | 0.542 | 0.00635 | ARHGEF7,CAMK2D |
| Neuropathic Pain Signaling In Dorsal Horn Neurons | 0.539 | 0.0099 | CAMK2D |
| Sumoylation Pathway | 0.531 | 0.00971 | ARHGDIA |
| IGF-1 Signaling | 0.524 | 0.00952 | PTK2 |
| Myelination Signaling Pathway | 0.52 | 0.00612 | ARF6,PTK2 |
| Role of Macrophages, Fibroblasts and Endothelial Cells in | 0.511 | 0.00602 | CAMK2D,DAAM1 |
| Role of MAPK Signaling in Promoting the Pathogenesis of | 0.495 | 0.00877 | ATP6V1E1 |
| Breast Cancer Regulation by Stathmin1 | 0.49 | 0.00505 | ARHGEF1,ARHGEF7,CAMK2D |
| Neuregulin Signaling | 0.486 | 0.00855 | ITGAV |
| Cholecystokinin/Gastrin-mediated Signaling | 0.48 | 0.0084 | PTK2 |
| Sphingosine-1-phosphate Signaling | 0.477 | 0.00833 | PTK2 |
| Renin-Angiotensin Signaling | 0.474 | 0.00826 | PTK2 |
| Neuroprotective Role of THOP1 in Alzheimer's Disease | 0.474 | 0.00826 | DPP4 |
| LXR/RXR Activation | 0.468 | 0.00813 | PCYOX1 |
| IL-15 Production | 0.468 | 0.00813 | PTK2 |
| Glioma Signaling | 0.462 | 0.008 | CAMK2D |
| FXR/RXR Activation | 0.459 | 0.00794 | PCYOX1 |
| Synaptic Long Term Potentiation | 0.443 | 0.00758 | CAMK2D |
| B Cell Receptor Signaling | 0.442 | 0.00472 | CAMK2D,PTK2,PTPRC |
| Atherosclerosis Signaling | 0.441 | 0.00752 | PCYOX1 |
| SNARE Signaling Pathway | 0.433 | 0.00735 | CAMK2D |
| Role of PKR in Interferon Induction and Antiviral Respons | 0.433 | 0.00735 | NPM1 |
| DHCR24 Signaling Pathway | 0.431 | 0.0073 | PCYOX1 |
| Iron homeostasis signaling pathway | 0.428 | 0.00725 | ATP6V1E1 |
| Adipogenesis pathway | 0.426 | 0.00719 | RBBP7 |
| MSP-RON Signaling In Cancer Cells Pathway | 0.423 | 0.00714 | PTK2 |
| Hereditary Breast Cancer Signaling | 0.418 | 0.00704 | NPM1 |
| Xenobiotic Metabolism General Signaling Pathway | 0.416 | 0.00699 | UGT1A6 |
| Endocannabinoid Cancer Inhibition Pathway | 0.407 | 0.0068 | PTK2 |
| Dilated Cardiomyopathy Signaling Pathway | 0.4 | 0.00667 | CAMK2D |
| Semaphorin Neuronal Repulsive Signaling Pathway | 0.4 | 0.00667 | ITGAV |
| Factors Promoting Cardiogenesis in Vertebrates | 0.393 | 0.00654 | CAMK2D |
| IL-10 Signaling | 0.391 | 0.00649 | BLVRA |
| Necroptosis Signaling Pathway | 0.387 | 0.00641 | CAMK2D |
| Aryl Hydrocarbon Receptor Signaling | 0.381 | 0.00629 | NEDD8 |
| Phagosome Formation | 0.38 | 0.00432 | ARF6,ITGAV,PTK2 |
| Hepatic Fibrosis Signaling Pathway | 0.378 | 0.00473 | ITGAV,PTK2 |
| Inhibition of ARE-Mediated mRNA Degradation Pathway | 0.374 | 0.00617 | EDC4 |
| HOTAIR Regulatory Pathway | 0.372 | 0.00613 | RBBP7 |
| HMGB1 Signaling | 0.365 | 0.00599 | RBBP7 |
| CXCR4 Signaling | 0.363 | 0.00595 | PTK2 |
| Germ Cell-Sertoli Cell Junction Signaling | 0.359 | 0.00588 | PTK2 |
| D-myo-inositol (1,4,5,6)-Tetrakisphosphate Biosynthesis | 0.341 | 0.00556 | PTPRC |
| D-myo-inositol (3,4,5,6)-tetrakisphosphate Biosynthesis | 0.341 | 0.00556 | PTPRC |
| FAK Signaling | 0.33 | 0.00384 | ARHGEF7,ITGAV,PTK2,SPARCL1 |
| MicroRNA Biogenesis Signaling Pathway | 0.329 | 0.00535 | ILF2 |
| Macrophage Classical Activation Signaling Pathway | 0.325 | 0.00529 | RBBP7 |
| NOD1/2 Signaling Pathway | 0.325 | 0.00529 | ARHGEF7 |
| Granulocyte Adhesion and Diapedesis | 0.325 | 0.00529 | PECAM1 |
| Production of Nitric Oxide and Reactive Oxygen Species i | 0.322 | 0.00524 | PCYOX1 |
| 3-phosphoinositide Degradation | 0.322 | 0.00524 | PTPRC |
| Xenobiotic Metabolism CAR Signaling Pathway | 0.32 | 0.00521 | UGT1A6 |
| Hepatic Fibrosis / Hepatic Stellate Cell Activation | 0.317 | 0.00515 | COL12A1 |
| D-myo-inositol-5-phosphate Metabolism | 0.314 | 0.0051 | PTPRC |
| Pulmonary Healing Signaling Pathway | 0.309 | 0.00503 | PECAM1 |
| Adrenomedullin signaling pathway | 0.309 | 0.00503 | PTK2 |
| Coronavirus Pathogenesis Pathway | 0.302 | 0.0049 | NPM1 |
| Sertoli Cell-Sertoli Cell Junction Signaling | 0.299 | 0.00485 | ITGAV |
| 3-phosphoinositide Biosynthesis | 0.299 | 0.00485 | PTPRC |
| HIF1α Signaling | 0.296 | 0.00481 | CAMK2D |
| Agranulocyte Adhesion and Diapedesis | 0.293 | 0.00476 | PECAM1 |
| ICOS-ICOSL Signaling in T Helper Cells | 0.291 | 0.00394 | CAMK2D,PTPRC |
| Calcium Signaling | 0.279 | 0.00455 | CAMK2D |
| Role of NFAT in Cardiac Hypertrophy | 0.274 | 0.00446 | CAMK2D |
| HER-2 Signaling in Breast Cancer | 0.27 | 0.00441 | ARF6 |
| Superpathway of Inositol Phosphate Compounds | 0.261 | 0.00427 | PTPRC |
| IL-12 Signaling and Production in Macrophages | 0.259 | 0.00424 | PCYOX1 |
| cAMP-mediated signaling | 0.259 | 0.00424 | CAMK2D |
| Role Of Osteoblasts In Rheumatoid Arthritis Signaling Pat | 0.25 | 0.0041 | CTSC |
| Wound Healing Signaling Pathway | 0.241 | 0.00397 | COL12A1 |
| Sperm Motility | 0.235 | 0.00389 | PTK2 |
| TEC Kinase Signaling | 0.234 | 0.00346 | ITGAV,PTK2 |
| PI3K Signaling in B Lymphocytes | 0.225 | 0.00338 | CAMK2D,PTPRC |
| Circadian Rhythm Signaling | 0.224 | 0.00373 | CAMK2D |
| Insulin Secretion Signaling Pathway | 0.22 | 0.00368 | CAMK2D |
| Protein Ubiquitination Pathway | 0.219 | 0.00366 | UBE2N |
| Opioid Signaling Pathway | 0.212 | 0.00357 | CAMK2D |
| Huntington's Disease Signaling | 0.209 | 0.00353 | HIP1 |
| CLEAR Signaling Pathway | 0.208 | 0.00351 | ATP6V1E1 |
| Pathogen Induced Cytokine Storm Signaling Pathway | 0 | 0.0027 | COL12A1 |
| Role Of Osteoclasts In Rheumatoid Arthritis Signaling Pat | 0 | 0.00324 | COL12A1 |
| Glucocorticoid Receptor Signaling | 0 | 0.00172 | KRT15 |
| Neutrophil Extracellular Trap Signaling Pathway | 0 | 0.00251 | COL12A1 |
| S100 Family Signaling Pathway | 0 | 0.0013 | CAMK2D |
| Chaperone Mediated Autophagy Signaling Pathway | 0 | 0.00311 | ATP6V1E1,GPX4 |
| RAR Activation | 0 | 0.00234 | CAMK2D |
| CD28 Signaling in T Helper Cells | 0 | 0.00193 | PTPRC |
| Lipid Antigen Presentation by CD1 | 0 | 0.00241 | ARF6 |
| CREB Signaling in Neurons | 0 | 0.00165 | CAMK2D |
| Systemic Lupus Erythematosus Signaling | 0 | 0.00187 | PRPF8,PTPRC |
| CDC42 Signaling | 0 | 0.00174 | ITGAV |
| Phospholipase C Signaling | 0 | 0.00268 | ARHGEF1,ARHGEF7,ITGAV |
| B Cell Development | 0 | 0.00204 | PTPRC |
| PKCθ Signaling in T Lymphocytes | 0 | 0.00179 | CAMK2D |
| IL-4 Signaling | 0 | 0.00173 | COL12A1 |
| Serotonin Receptor Signaling | 0 | 0.00213 | CAMK2D |
| NF-κB Signaling | 0 | 0.00175 | UBE2N |
| T Cell Receptor Signaling | 0 | 0.00162 | PTPRC |
| G-Protein Coupled Receptor Signaling | 0 | 0.00284 | CAMK2D,PTK2 |
| Systemic Lupus Erythematosus In T Cell Signaling Pathw | 0 | 0.00156 | PTK2 |

**Figure 5.**
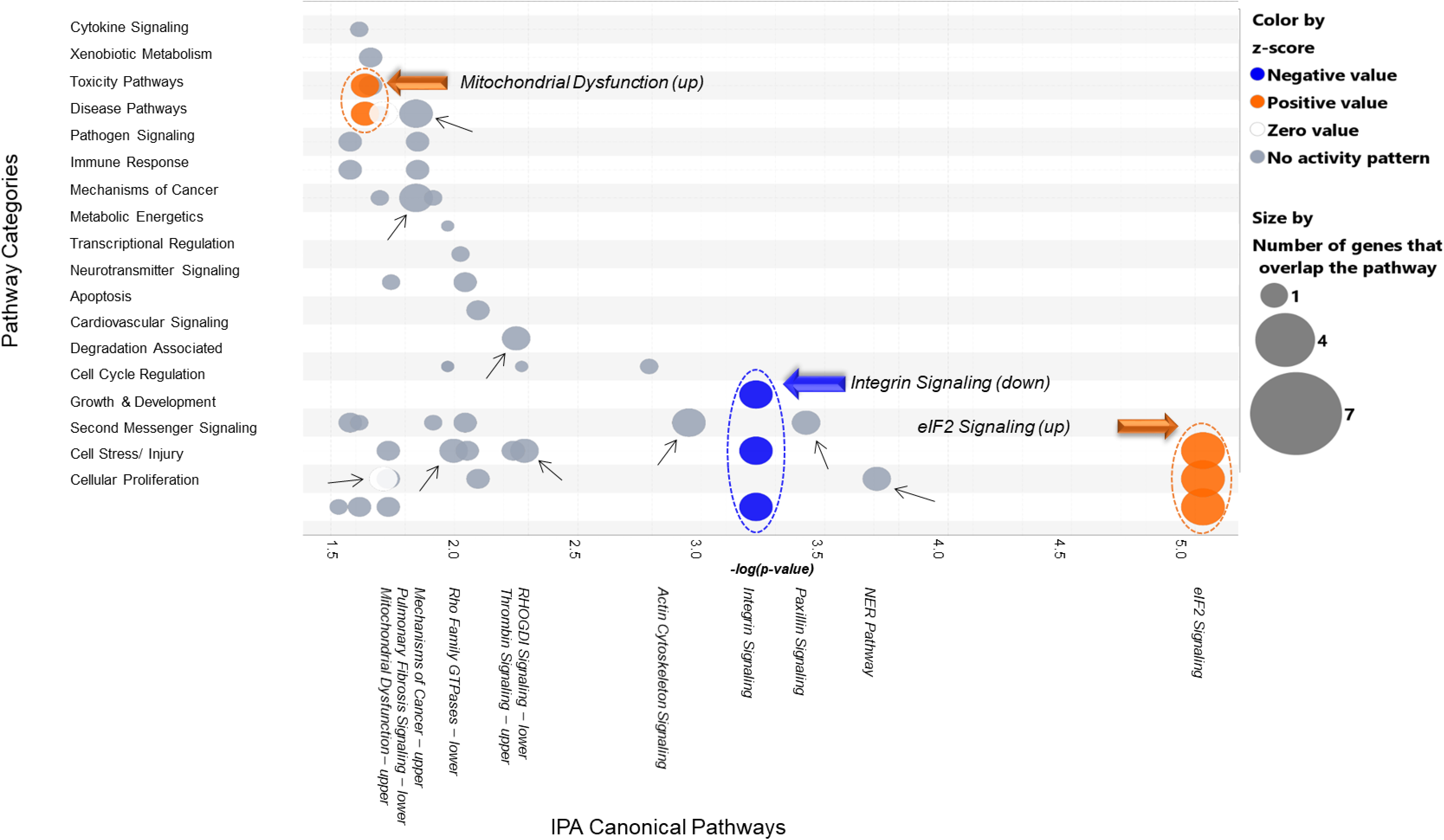
2D Bubble Chart of Top Pathways Attenuated/Reversed by hHPX post Cl_2_ Exposure. This 2D bubble chart illustrates pathway “categories” (y-axis) vs. canonical pathways (x-axis; negative log of the right-tailed Fisher’s Exact Test’s p-value) associated with the proteins that were found to be either attenuated or reversed by the presence of human hemopexin in mouse lungs following Cl2 exposure. The small (line arrows) and larger (colored arrows) are pointing toward “key” pathways found to be associated with each pathway category. The orange and blue color codes are highlighting the predicted pathway activities as either increased or decreased respectively, which is indicated by there calculated z-scores. The white and grey color codes are associated with either an unclear or balanced mix of activities based on the proteins identified, or simply no predicted alteration in pathway activities, which doesn’t take away from the significant associations between the proteins of interest and their **respective pathways, which is left to be determined via validation regardless.**

### Biochemical validation of proteomics data

Western blotting studies using antibodies against elF2a and phosphorylated elF2a showed that at 24 h post exposure to Cl_2_ there was a large increase of the phosphorylated form of elF2a in the lungs of vehicle but not hHPX treated mice (**Figures 6A, B**). Total elF2a amounts remained unchanged. An increase of phosphorylation indicates activation of elF2a by one of the four kinases and results in an overall decrease of protein transcription and translation (48). These data are consistent with the proteomics and systems biology conclusions mentioned above. On the other hand there eLF2a was not phosphorylated in the lung of mice at 6 h post exposure (**Figure 6C**), at a time that heme plasma levels were at not different from their air control value (**Figure 1B**).

**Figure 6.**
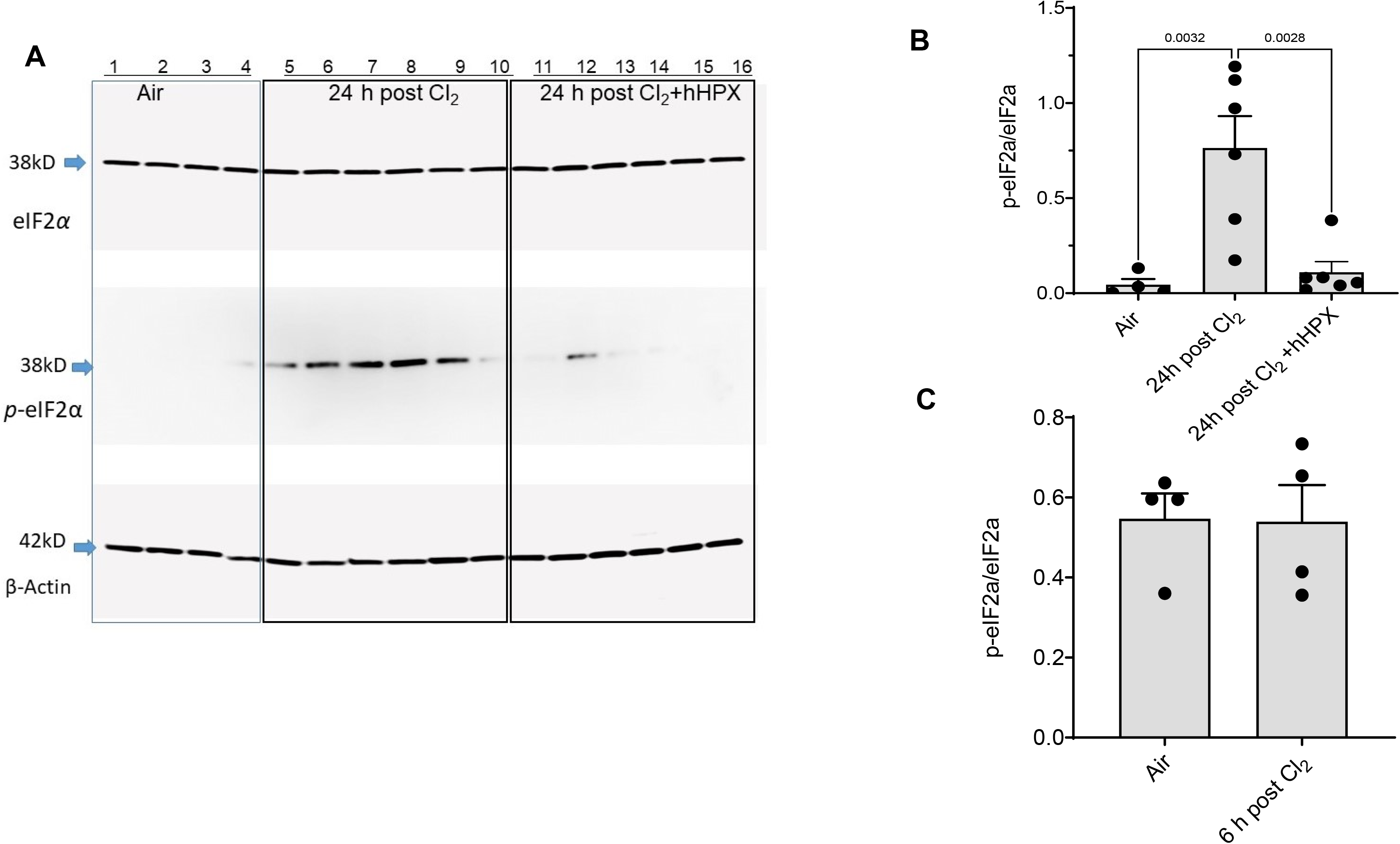
Western blotting studies of eIF2a in lung tissues post Cl_2_ exposure. Lung tissues of mice exposed to air or Cl2 (500 ppm for 30 min) and injected with hHPX (5 µg/g BW) at 1 h post exposure and sacrificed at 24 h later were probed with antibodies against elF2a or its phosphorylated form and actin as described in the METHODS. (**A**) Western blots. Each lane represents results from a different mouse. (**B and C**). Quantitation of of the ratio of phosphor-elF2a/total elF2a in the lungs of mice at 24 and 6 h post exposure to Cl_2_. Individual points and means ± 1 SEM; statistical analysis by one way analysis of variance followed by the Dunn–Šidák test for multiple comparisons (GraphPad Prism 9).

### Generation of recombinant hemopexin in plants

We have generated two forms of recombinant hemopexin in *Nicotiana benthamiana* plants. Both products, standard recombinant plant-based HPX (Std-prhHPX) and Gal-prhHPX contained high content of glycans with multiple mannose residues (M6-M9) as well as forms containing plant-specific glycans. The second product (Gal-prhHPX) was partially galactosylated, but far less so than hHPX and contained high levels of multi-mannose species and plant-specific glycans. Neither product was sialylated. Their pharmacokinetics of these two products in the plasma of air and Cl_2_ exposed mice following injection of 5 µg/g BW are shown in **Figure 7A**. Gal-prhHPX achieved much higher concentrations in the plasma as compared with Std-prhHPX, most likely due to improved glycosylation. However, the peak concentration of Gal-prhHPX and its ACU24 value were considerably lower than of hHPX, (2800 vs. 9500 ng/ml and 23.29 vs. 174.6 hr*mg/L).

**Figure 7.**
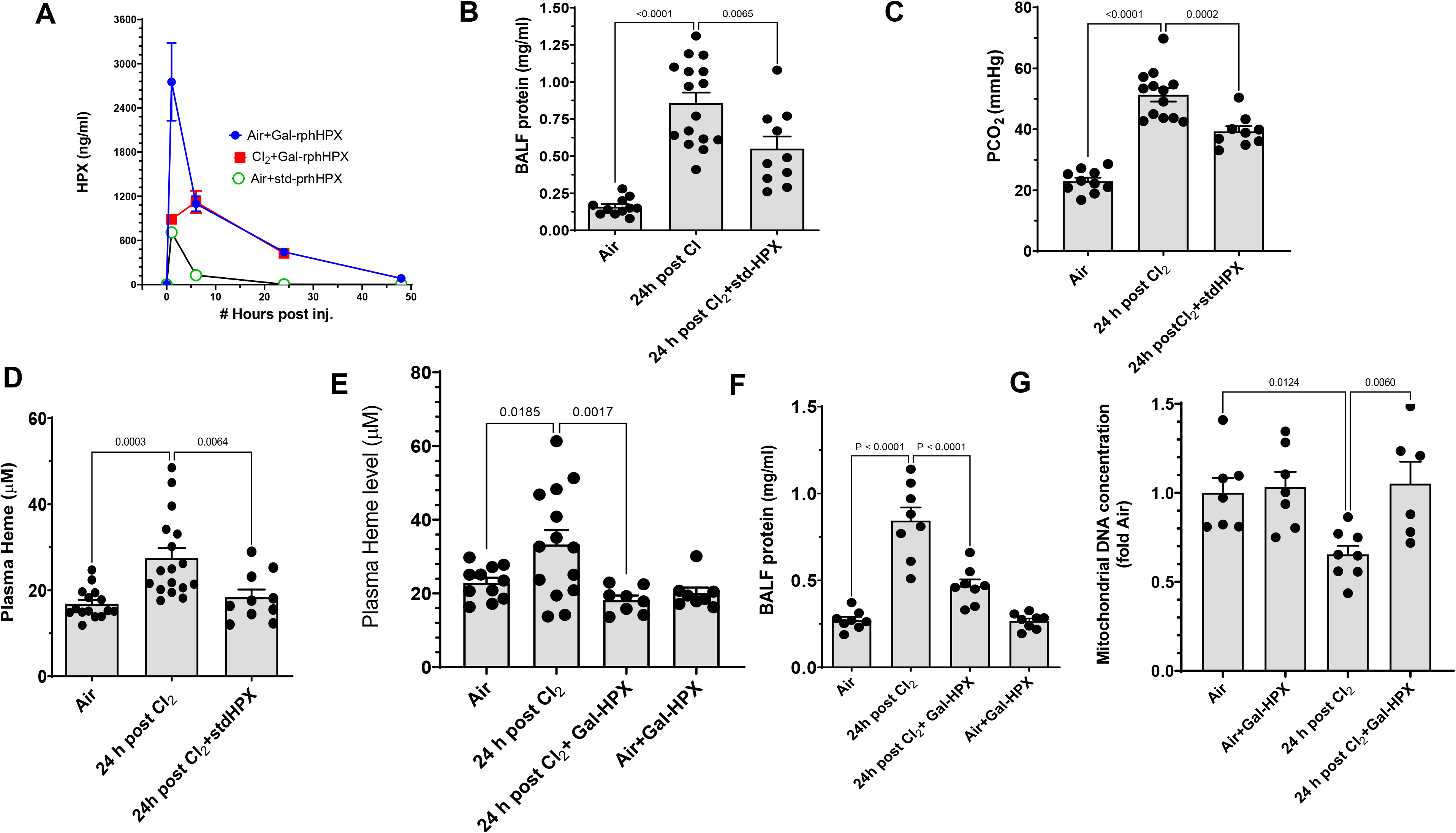
(**A**). **Pharmacokinetics of recombinant hemopexin.** C57BL/6 mice were exposed to air or Cl_2_ (500 ppm for 30 min). One h post exposure they were injected intramuscularly with either Gal-prhHPX or Std-prhHPX (hHPX; 5 µg/g BW). Mice were then sacrificed at the indicated intervals and the concentration of hHPX in their plasmas was measured by ELISA as described in the methods. Each symbol represents a different mouse. Lines connect values from different mice obtained at the same time. **Recombinant hemopexin decreases heme and acute lung injury.** C57BL/6 mice were exposed to air or Cl_2_ (500 ppm for 30 min) and returned to room air. One h post exposure they were injected intramuscularly with either std-prhHPX (10 µg/g BW) (**panels B, C, D**) or Gal-prhHPX (5 µg/g BW) (**E, F, G**). Mice were then sacrificed at 24 h post exposure, their lungs were lavaged and the indicated variables were measured. Each point represents a different animal. Means ± 1 SEM; statistical analysis by one way analysis of variance followed by the Dunn–Šidák t-test for multiple comparisons.

### Recombinant hemopexin decreases Cl_2_-induced acute lung injury

In our next series of experiments, we tested the efficacy of the two forms of recombinant hemopexin (Std-prhHPX and Gal-prhHPX) to decrease acute lung injury when injected intramuscularly 1h post Cl_2_. For Std-prhHPX we opted to inject 10 µg/g BW to partially offset the known rapid clearance in the plasma (**Figure 7**). As seen in **Figure 7 B,C,D** injection of Std-prhHPX decreased heme levels to the air values and decreased the concentration of proteins in the BAL. In addition, it decreased arterial P_CO2_ which was markedly elevated after exposure to Cl_2_. Injection of 5 µg/g BW of Gal-prhHPX decreased heme levels (**Figure 7E**), protein in the BALF (Figure 8F) and reversed injury to mitochondrial (**Figure 7G**). Neither Std-prhHPX nor Gal-prhHPX had any effect on any of these variables in air breathing mice.

## Discussion

The major novel findings of this study are: (1) exposure of adult male and female mice to Cl_2_, in concentrations likely to be encountered within one mile of the epicenter of industrial accidents, result in increased concentrations of free heme in the plasma, damage to mtDNA, the development of lung injury and major changes in the lung proteome; (2) post Cl_2_ exposure injection of hHPX, the most efficient scavenger of free heme, decreases the severity of lung and mtDNA injury, mitigates proteomic changes and increases survival and (3) two recombinant plant based forms of hHPX mimic the effects of hHPX, in spite of the fact that they lacked the glycosylation profiles of hHPX and were cleared faster from the plasma.

Heme is maintained at low levels(10) by serum albumin, haptoglobin and hemopexin (16), a plasma protein with the highest known binding affinity to free heme (K_d_ near 10^−13^ M). The heme–hemopexin complex is transported to liver cells where it binds to CD91/LRP1, is internalized via receptor-mediated endocytosis and metabolized to biliverdin, carbon monoxide and iron by heme oxygenases (HO)(7, 8). We have shown that a single intramuscular injection of hHPX, in male and female C57BL/6 or humanized mice with sickle cell disease (SCD) post Cl_2_ and Br_2_ exposure decreases acute and chronic lung injury and improves survival(3, 5, 6, 8). Furthermore, HO-1^(−/−)^ mice are more susceptible to halogen-induced injury(5, 7). hHPX has also been shown to decrease cardiopulmonary and systemic injury in a number of hemolytic diseases(19, 24, 29, 30, 54). Our pharmacokinetic measurements showing decreased levels of both hHPX and prhHPX in the plasma of mice post exposure to Cl_2_, when large levels of free heme are present, agree with the study of Poillerat *et al*.(38) who demonstrated that the pharmacokinetics of human hemopexin are altered significantly in a mouse model of hemolytic anemia, when large amounts of free heme are present in the plasma. However, our data indicates that hemopexin regulates the activity of a large number of additional proteins as was the case in air breathing mice (supplemental figures S1).

hHPX reversed to various degrees the upregulation of a number of proteins involved in the inflammatory process (such as Platelet endothelial cell adhesion molecule, focal adhesion kinase 1, Rho guanine nucleotide exchange factor 1) and modulated the expression of a number of proteins involved in mitochondrial function. Among the most important proteins reversed by hemopexin were Glutathione Peroxidase 4, a major antioxidant involved in the reduction of complex hydroperoxides using reduced glutathione the suppression of which leads to ferroptosis(46, 53), acetyl-CoA synthetase 2 (AceCS2), a mitochondrial enzyme which plays a significant role in acetate oxidation needed to generate ATP and heat (40), ATP-dependent RNA helicase A which plays a pivotal role in mRNA translation initiation process (22) and others. On the opposite side of the spectrum, exposure to Cl_2_ resulted in upregulation of Platelet endothelial cell adhesion molecule which plays an important role in the transmigration of monocytes, neutrophils and NK cells(50), integrin alpha-3, which is part of the integrin complex that regulates cytokine activation and plays an important role in inflammation(14). Cathepsin C,(49), involved in the activation of elastase and cathepsin G in neutrophils which promote tissue damage in addition to microbial killing (35), histone H2B type 2-B, which plays an important role in DNA repair (39) and others. The demonstration that prhHPX can interact and remove heme from the plasma is highly significant since hHPX is isolated from human plasma. Thus, it is costly and may not be readily available in quantities to conduct studies in larger animals and Phase I clinical trials. Plant-based production systems for biopharmaceutical proteins are attractive alternatives to mammalian cell, yeast, or bacterial systems(12). Additional benefits include low cost, scalability, long term stability, lack of non-animal components, and rapid deployment of new therapies.

Two of the most striking findings of our systems biology analysis were the demonstrations that injection of hHPX upregulated the Cl_2_ induced downregulation of eIF2 signaling and reversed mtDNA damage. These findings were validated by western blotting studies showing that injection of hHPX reversed or prevented the phosphorylation of eIF2a, which is linked to its activation, in lung tissue and by measurements of mtDNA concentrations by RT-PCR. Previous studies have shown that phosphorylation of eLF2a at serine 51 results in decreased translation of a number of eukaryotic proteins and favors the synthesis of activating transcription factor 4 (ATF4) which translocates to the nucleus and initiates transcription of proteins responsible for the removal of misfolded proteins (9, 48) and initiation of autophagy (11, 18). elF2a phosphorylation also contributes to the activation of NF-κB and production of inflammatory cytokines following viral infections(43). Phosphorylation of elF2a is regulated by four kinases (general control nonderepressible 2 [GCN2], heme-regulated inhibitor [HRI], protein kinase RNA-like endoplasmic reticulum kinase [PERK], and protein kinase R [PKR]), which are in turn activated by, oxidative stress (48). We have previously shown that exposure of BALB/c female mice, in which the fur was removed, to Cl_2_ (400 ppm for 30 min) led to phosphorylation of PERK in both the skin and lung tissues, phosphorylation of eIF2a in the lung and increased levels of both CHOP and ATF6 in lung and skin and increased transcript of IL-1b, IL6 and TNFa in lung tissue at 6 h post exposure(28). These were the first data to indicate the onset of UPR response post exposure to Cl_2_ gas, resulting from increased oxidant stress (27, 51, 52) The UPR response was found to contribute to the development of pulmonary fibrosis and emphysema-type disease at 14-21 days post exposure of mice to the halogen bromine and post exposure administration of hHPX reversed these changes (3). However, our data provide the first demonstration that hemopexin prevents the phosphorylation of eIF2a in Cl_2_ exposed mice and modulates inflammatory and non-inflammatory biological processes by modifying the lung proteome. Whether this is also true for prhHPX has to be determined.

Oxidative stress damages the sugar-phosphate backbone causing strand breaks in the mitochondrial DNA (mtDNA) (42). Mitochondrial DNA loss has been observed after exposure to ethanol (31), lipopolysaccharide (44, 45), Pseudomonas Aeruginosa lung injury (23), ventilator induced lung injury (15), and oxidant induced injury to the pulmonary endothelium (13).

In previous studies using redox sensitive dyes, as well as electron spin resonance, we showed conclusively that exposure of rat alveolar type II cells in primary culture and H441 cells, a human club-cell line, to Cl_2_ *in vitro*, increases the steady state concentrations of reactive intermediates in their cytoplasm and mitochondria. Mitochondrial reactive species were detected even 24 h post exposure (20, 25). Increased concentrations of reactive species, quantified as increased lung tissue lipid peroxidation, carbonylation and F2-isoprostanes formation, a group of prostaglandin F2-like compounds derived from the nonenzymatic oxidation of arachidonic acid by Cl_2_ or HOCl, its hydration product, were also detected in the lungs of mice exposed to either Cl_2_ or Br_2_ and returned to room air for 6-24 h (3, 6, 52). We also found that lung epithelial cells exposed to Cl_2_ and returned to room air for 6 h, developed compromised mitochondrial bioenergetics, as shown by decreases in maximal Oxygen Consumption Rate (OCR), Reserve Capacity and mitochondrial membrane potential, and a concomitant increase of non-mitochondrial OCR, an index of reactive species formation. Treatment of cells with the mitochondrial redox modulator MitoQ, attenuated these bioenergetic defects and decreased mitochondrial reactive species (20). These findings highlight the importance of Cl_2_ induced injury to mitochondria. This is the first study to report mtDNA injury post exposure to Cl_2_ in vivo.

In summary, our proteomics, biochemical and physiological studies provide new insight into the mechanisms by which exposure to Cl2 gas damages the pulmonary system and highlight the potential of human and recombinant hemopexin in reversing lung injury and improve mortality.

## Acknowledgments

These studies were supported by R21 ES032956-01 from NIEHS, 2R01HL031197-28 from NHLBI and a grant from the Center for Clinical and Translational Science of the University of Alabama at Birmingham to SM. TJ was supported by 5R2 ES031559 from NIEHS; ASA was supported by 1R21ES034226; JM and KK were supported by the Mass Spectrometry/Proteomics Shared Facility 3P30CA013148 from the NCI.

## SUPPLEMENTAL MATERIAL

### Supplementary Figure Legends

**Supplement Figure 1.**
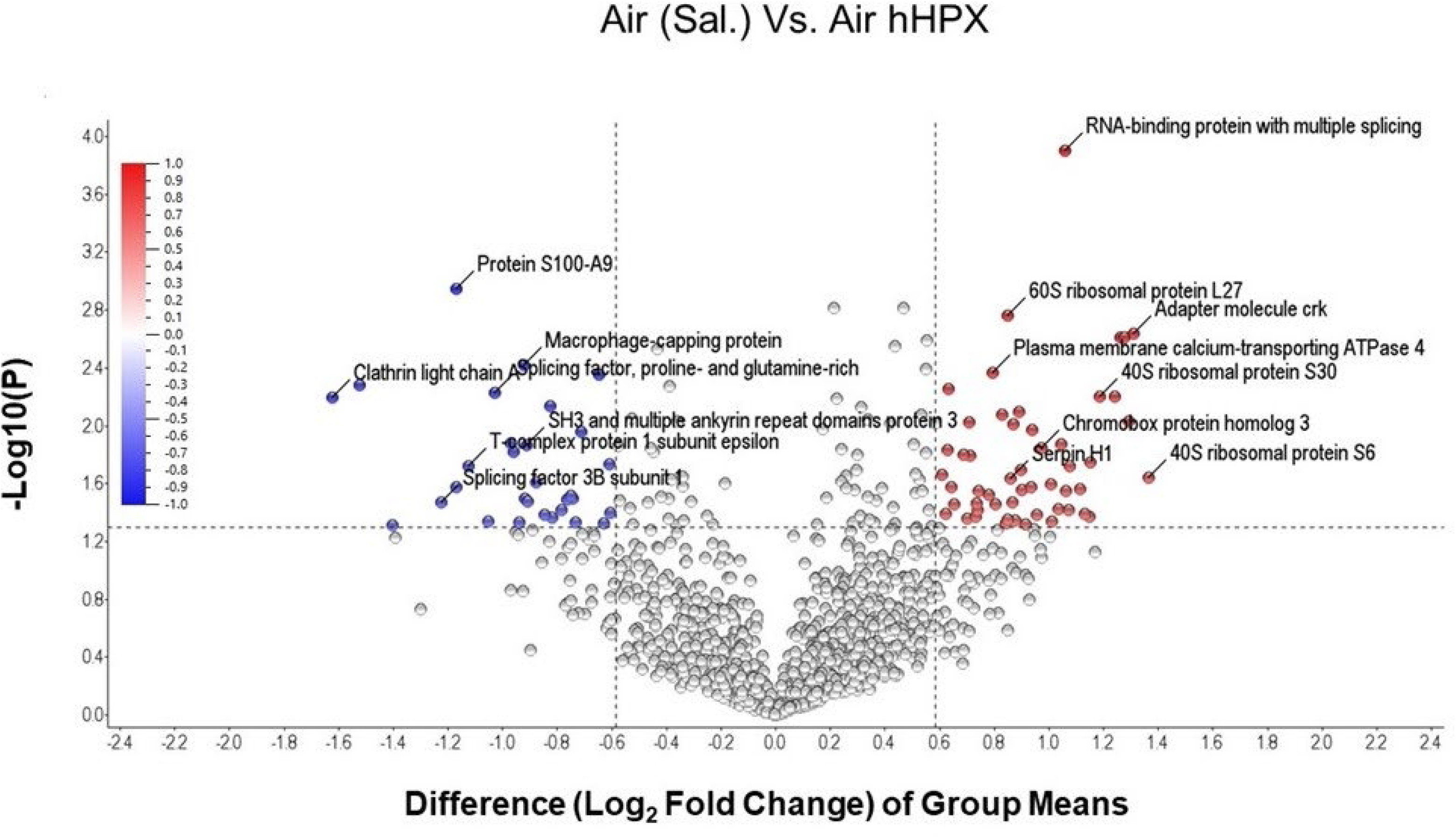
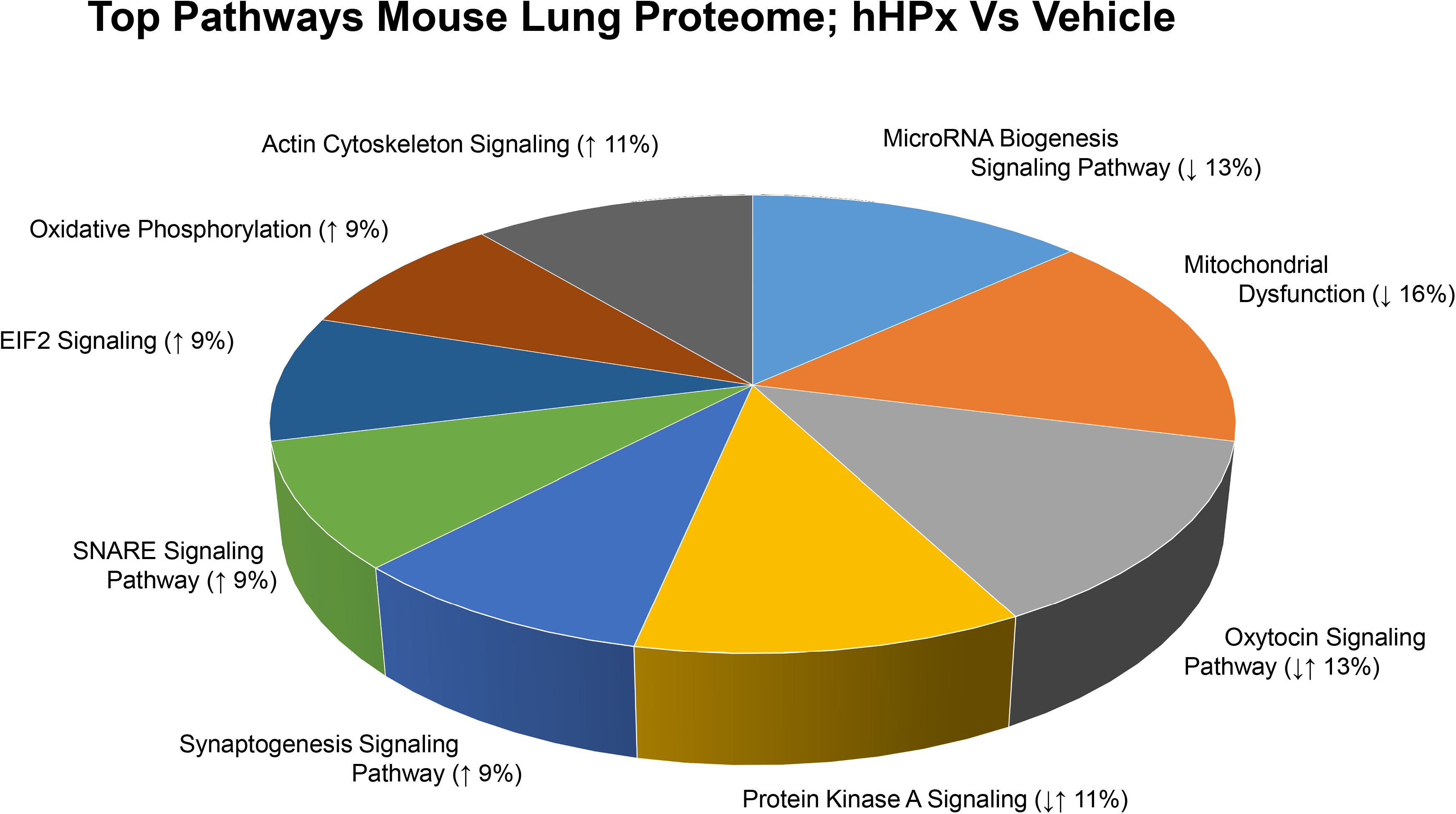
A. Global protein changes in the lungs of C57BL/6 mice exposed to air and injected with saline or human hemopexin. C57BL/6 mice were exposed to air and received a single dose of human hemopexin (5 µg/ g BW) or an equal volume of sterile saline intramuscularly. Twenty four hours later the mice were sacrificed and their lungs were removed and proteins were processed for global proteomics analysis as discussed in the METHODS. Volcano plot, of the Log10 p-value vs. Log2 fold change (Air/Saline vs. Air/hHPX) demonstrating the distribution of the entire data set of proteins with upper limits (above the line) indicating statistically significant changes, and outer limits (to the right and left of each line) indicating significant fold changes as outlined in the methods section under statistics. Note that while fold change is visualized as Log2, the cut-off value of ± 1.5 were applied to the fold change prior to logging, thereby yielding the indicated ±0.6 limits. **B.** System analysis utilizing the top-95 statistically significant proteins with fold change ≥±1.5 () allowed us to categorize them according to: (a) cellular locations and (b) biological processes. The number of proteins associated with each location or process were summed and normalized to 100 (of note, each protein can be associated with more than one location or process). The resultant pie charts are indicative of the normalized percent proteins associated with each category within cellular localizations and biological functions. We have added an asterisk next all segments that we are particularly interested in for each pie chart, these cellular locations and biological functions are discussed in the text.

